# A Universal Bacteriophage T4 Nanoparticle Platform to Design Multiplex SARS-CoV-2 Vaccine Candidates by CRISPR Engineering

**DOI:** 10.1101/2021.01.19.427310

**Authors:** Jingen Zhu, Neeti Ananthaswamy, Swati Jain, Himanshu Batra, Wei-Chun Tang, Douglass A. Lewry, Michael L. Richards, Sunil A. David, Paul B. Kilgore, Jian Sha, Aleksandra Drelich, Chien-Te K. Tseng, Ashok K. Chopra, Venigalla B. Rao

## Abstract

A “universal” vaccine design platform that can rapidly generate multiplex vaccine candidates is critically needed to control future pandemics. Here, using SARS-CoV-2 pandemic virus as a model, we have developed such a platform by CRISPR engineering of bacteriophage T4. A pipeline of vaccine candidates were engineered by incorporating various viral components into appropriate compartments of phage nanoparticle structure. These include: expressible spike genes in genome, spike and envelope epitopes as surface decorations, and nucleocapsid proteins in packaged core. Phage decorated with spike trimers is found to be the most potent vaccine candidate in mouse and rabbit models. Without any adjuvant, this vaccine stimulated robust immune responses, both T_H_1 and T_H_2 IgG subclasses, blocked virus-receptor interactions, neutralized viral infection, and conferred complete protection against viral challenge. This new type of nanovaccine design framework might allow rapid deployment of effective phage-based vaccines against any emerging pathogen in the future.

## Introduction

Rapid discovery of safe and effective vaccines against emerging and pandemic pathogens such as the novel coronavirus SARS-CoV-2^1, 2^ requires a “universal” vaccine design platform that can be adapted to any infectious agent^3, 4^. It should allow incorporation of diverse targets, DNAs, and proteins (multicomponent), full-length proteins as well as peptides and domains, in various combinations (multivalent). Such a multiplex platform would not only compress the timeline for vaccine discovery but also offers critical choices for selecting the most effective vaccine candidate(s) without going through iterative design cycles^4–6^.

Numerous SARS-CoV-2 vaccine candidates have been developed at record-breaking pace to quell this devastating pandemic and many are in clinical trials^5, 7–11^. Two of these, both mRNA based, have been approved by FDA for emergency use authorization. However, innovative platforms are still desperately needed because future pathogens might be more complex and their vaccine targets may not be as well-defined^3, 4, 11^. A drawback of the current platforms is that they are largely limited to a single vaccine target. They also lack sufficient engineering flexibility to generate multiplex vaccines, require strong chemical adjuvants to boost immune responses, and may not be accessible to resource-poor countries^4–6, 12^. Here, we present a “universal” nanovaccine platform by CRISPR engineering of bacteriophage (phage) T4 that can rapidly generate multiplex vaccine candidates against any emerging pathogen during epidemic or pandemic situations.

Tailed bacteriophages such as T4 are the most abundant and widely distributed organisms on Earth. T4 belongs to *myoviridae* family, infects *Escherichia coli*, and has served as an extraordinary model organism in molecular biology and biotechnology^13^. It consists of a 1200 Å long and 860 Å wide prolate head (or capsid) that encapsidates ~170 kb linear DNA genome, and a ~1400 Å long contractile tail with six long tail fibers emanating from a baseplate present at the tip of the tail^14^ (Figure 1). The head, the principal component for vaccine design, is assembled with 155 hexameric capsomers of the major capsid protein gp23* (“*” represents cleaved mature form), 11 pentamers of gp24* at eleven of twelve vertices, and 1 dodecameric portal protein gp20 at the unique twelfth vertex^15, 16^ (Figure 1g).

**Fig. 1.**
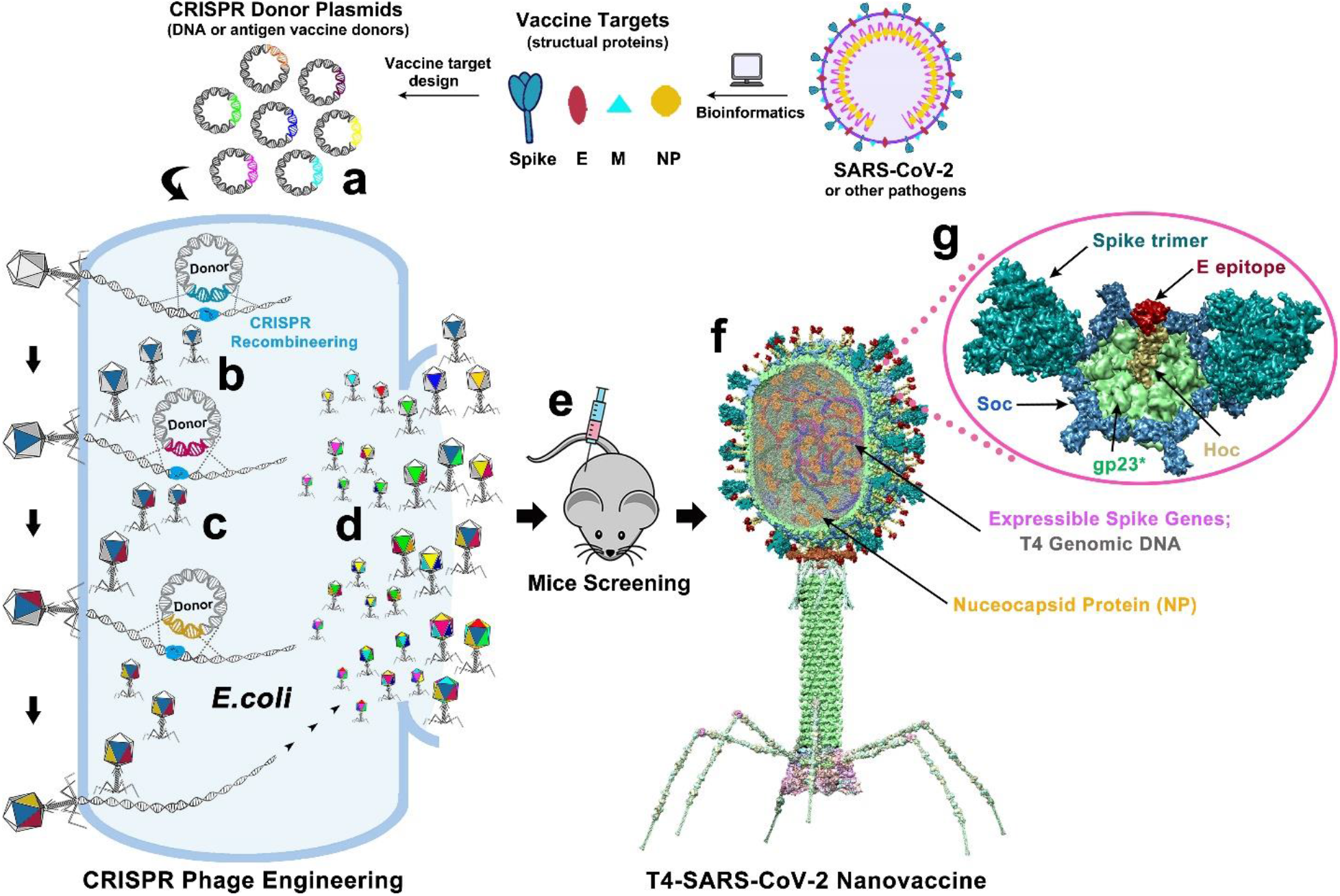
Design of T4-SARS-CoV-2 nanovaccine by CRISPR engineering. Engineered DNAs corresponding to various components of SARS-CoV-2 virion are incorporated into bacteriophage T4 genome. Each DNA was introduced into *E. coli* as a donor plasmid **(a)**, recombined into injected phage genome through CRISPR-targeted genome editing **(b)**. Different combinations of CoV-2 inserts were then generated by simple phage infections and identifying the recombinant phages in the progeny **(c)**. For example, recombinant phage containing CoV-2 insert #1 (dark blue) can be used to infect CRISPR *E. coli* containing Co-V2 insert containing donor plasmid #2 (dark red). The progeny plaques obtained will contain recombinant phage #3 with both inserts #1 and #2 (dark blue plus dark red) in the same genome. This process was repeated to rapidly construct a pipeline of multiplex T4-SARS-CoV-2 vaccine phages **(d)**. Selected vaccine candidates were then screened in a mouse model **(e)** to identify the most potent vaccine **(f)**. Structural model of T4-SARS-CoV-2 Nanovaccine showing an enlarged view of a single hexameric capsomer **(g)**. The capsomer shows six subunits of major capsid protein gp23* (green), trimers of Soc (blue), and a Hoc fiber (yellow) at the center of capsomer. The expressible spike genes are inserted into phage genome, the 12 aa E external peptide (red) is displayed at the tip of Hoc fiber, S-trimers (cyan) are attached to Soc subunits, and nucleocapsid proteins (yellow) are packaged in genome core. See Results, Materials and Methods, and Supplementary Video for additional details.

The T4 capsid is coated with two nonessential proteins; Soc (<u>*s*</u>mall <u>*o*</u>uter <u>*c*</u>apsid protein) (9.1 kDa; 870 copies per capsid) and Hoc (<u>*h*</u>ighly antigenic <u>*o*</u>uter <u>*c*</u>apsid protein) (40.4 kDa; 155 copies per capsid)^16, 17^ (Figure 1g). Soc is a trimer, bound to quasi three-fold axes, and acts as a “molecular clamp” by clasping adjacent capsomers^18^. Hoc is a 170 Å-long fiber containing a string of four Ig-like domains with its N-terminal domain exposed at the tip of the fiber^19^. Soc reinforces an already stable T4 capsid while Hoc helps phage to adhere to host surfaces^20^.

The above constitutes an ideal architecture to develop a universal vaccine design template. Further, our four decades of genetic, biochemical, and structural studies on phage T4 including the recently developed CRISPR (<u>c</u>lustered <u>r</u>egularly <u>i</u>nterspaced <u>s</u>hort <u>p</u>alindromic <u>r</u>epeats) phage engineering^21, 22^ provide an extraordinary resource. The atomic structures of all the capsid proteins including Soc, Hoc, as well as the entire capsid have been determined^15, 18, 19, 23, 24^. Soc and Hoc can be used as efficient adapters to tether foreign proteins to T4 capsid^25, 26^. Both have nanomolar affinity and exquisite specificity, allowing up to ~1,025 molecules of full-length proteins, domains, and peptides to be arrayed on capsid^27, 28^. T4 capsids so decorated with pathogen epitopes mimic PAMPs (pathogen-associated molecular patterns) of natural viruses and stimulate strong innate as well as adaptive immune responses^29^.

To develop a universal vaccine design template, we took advantage of a large amount of nonessential genetic space available in T4 genome. Using SARS-CoV-2 as a model pathogen, we inserted a number of viral components into phage including spike (S)^30, 31^, envelope (E)^32^, and nucleocapsid proteins (NP)^2, 33^ by CRISPR engineering as DNA and/or protein (Figure 1a-d; Video). These were then combined by simple phage infections to create a collection of recombinant phages containing different combinations of vaccine targets. In a few weeks, a pipeline of vaccine candidates in dozens of combinations were generated, demonstrating the unprecedented engineering power and flexibility of this approach. When tested in mouse and rabbit models, one of the candidates, T4 phage decorated with S-trimers (Figure 1e-g; Video) elicited robust immunogenicity and ACE-2 (angiotensin converting enzyme-2) receptor^34^ blocking and virus neutralizing antibodies that conferred complete protection against virus challenge in a mouse model.

Our studies thus established a “blueprint” for a new type of nanovaccine framework for rapid and multiplex design of effective vaccine candidates that can potentially be applied to any emerging pathogen in the future.

## Results

### Construction of T4-SARS-CoV-2 recombinant phages by CRISPR engineering

A series of CRISPR-*E. coli* strains were constructed by inserting SARS-CoV-2 gene segments into T4 phage genome. Each strain harbored two plasmids (Fig. 2a); a “spacer” plasmid expressing the genome-editing nuclease, either type II Cas9 or type V Cpf1, and CRISPR RNAs (crRNAs or “spacer” RNAs) corresponding to target protospacer sequence(s) in phage genome, and a second “donor” plasmid containing the SARS-CoV-2 sequence. The latter also has ~500 bp homologous flanking arms of phage genome corresponding to the point of insertion. Four nonessential regions of the genome were chosen for insertion of various SARS-CoV-2 genes (Fig. 2b, I-IV). When these *E. coli* were infected by T4, a double-stranded break would occur in the protospacer sequence by Cas9 or Cpf1 that inactivates the phage genome and no phage should be produced. However, the highly recombinogenic T4 phage allows efficient recombination between the cleaved DNA and the donor plasmid through the flanking homologous arms, transferring the CoV-2 gene into phage genome and propagating it as part of phage infection (Fig. 2a). The same strategy was used to introduce many other genetic modifications including deletions by simply creating that modification in the donor plasmid and various modifications were combined as desired by simple phage infections of appropriate CRISPR *E. coli* (Fig. 1).

**Fig. 2.**
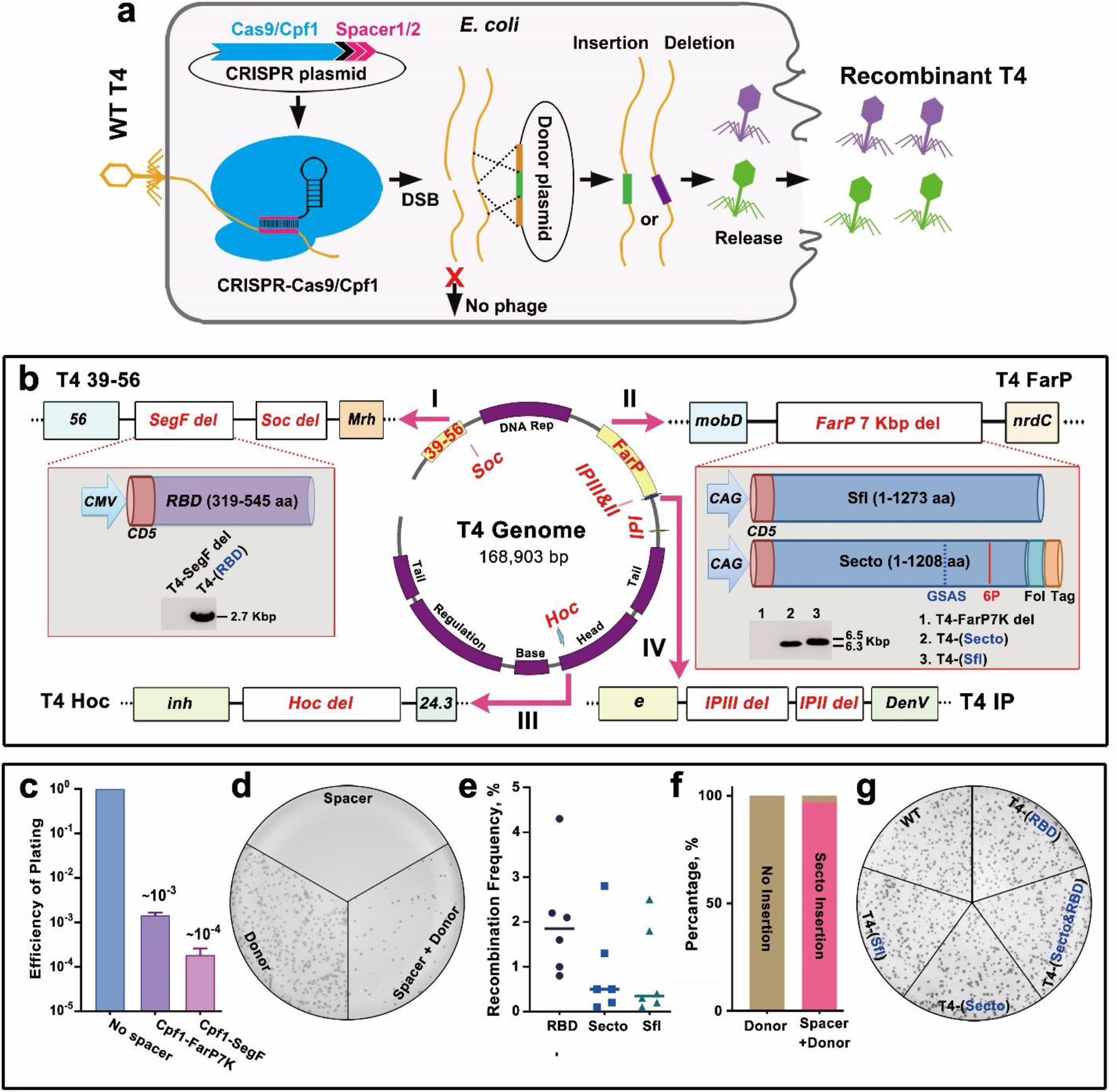
Construction of T4-SARS-CoV-2 recombinant phages by CRISPR engineering. **a.** Schematic of T4 CRISPR engineering. **b.** Four nonessential regions of T4 genome are chosen for deletion and insertion of various SARS-CoV-2 genes (shown in red; SegF/Soc, FarP, IP, and Hoc). 6P, six proline substitutions in S-ecto (F817P, A892P, A899P, A942P, K986P, and V987P). Fol, T4 fibritin motif Foldon for efficient trimerization. Tag, octa-histidine and twin-strep tags. Furin cleavage site RRAR was mutated to GSAS to stabilize trimers in a prefusion state^31^. **c.** Efficiency of plating (EOP) of representative Cpf1-FarP7K and Cpf1-SegF spacers. **d.** Plate showing plaques from phage infection of bacteria containing Cpf1-FarP7K spacer only, S-ecto donor only, or Cpf1-FarP7K spacer plus S-ecto donor. **e.** Recombination frequency of three spike gene (RBD, S-ecto, and S-fl) insertions. **f.** DNA sequencing of thirty independent plaques showed that >95% of the plaques generated in S-ecto recombination contained the correct S-ecto insert. **g.** Plate showing that the wild-type (WT), T4-RBD, T4-S-fl, T4-S-ecto, and T4-(S-ecto)-RBD recombinant phages had similar plaque size.

Initially, we constructed “acceptor” phages by deleting certain known nonessential segments of the phage genome^35^; ~18 kb FarP, ~11-kb 39-56, or both (~29 kb), that created space for CoV-2 insertions (Supplementary Fig. 1a,b). But the yields of these phages were low, ~1-2 orders of magnitude lower than the wild-type (WT) phage (Supplementary Fig. 1a,b). Since yield is critical for vaccine manufacture, we then constructed shorter deletions in which ~700 bp SegF within 39-56 and ~7 kb segment within FarP were deleted (Fig. 2b I, II). The yields of these phages (*7del.SegFdel.*T4) were similar to the WT phage, suitable for SARS-CoV-2 vaccine design.

Next, three spike gene variants^31^ corresponding to i) 1273-amino acid (aa) WT full-length (S-fl), ii) 1208 aa ectodomain (S-ecto, aa 1-1208), and iii) 227 aa receptor binding domain (RBD, aa 319-545) were engineered as expressible cassettes and inserted into *7del.SegFdel.* T4 (Fig. 2b, Supplementary Fig. 1c). The spike genes were codon-optimized and kept under the control of a strong mammalian expression promoter, either CMV or CAG, and a human CD5 signal peptide fused to the N-terminus for efficient secretion (Fig. 2b I, II, Supplementary Fig. 1d). The S-ectodomain recombinant contained additional mutations including six proline substitutions that imparted greater stability and ~10-fold greater expression, as was described by Hsieh *et al*.^31^ (Fig. 2b, Supplementary Fig. 1d).

Control CRISPR *E. coli* containing Cas9/Cpf1-spacer plasmid but lacking the spike gene donor plasmids yielded very few or no plaques when infected with *7del.SegFdel* phage (plating efficiency, <10^−4^ to 10^−3^; Fig. 2c,d, Supplementary Fig. 1e) whereas those containing both the plasmids produced plaques at a recombination frequency of up to ~4.5% (Fig. 2d,e). DNA sequencing confirmed that >95% of these plaques were true recombinants with correct insert (Fig. 2f) and showed similar plaque forming ability as the WT phage (Fig. 2g). A similar CRISPR strategy was used for creating deletions and/or insertions at the other sites [internal protein (IP)II, IPIII, Hoc, and Soc] (Fig. 2b; Supplementary Fig. 1f).

**Supplementary Fig. 1.**
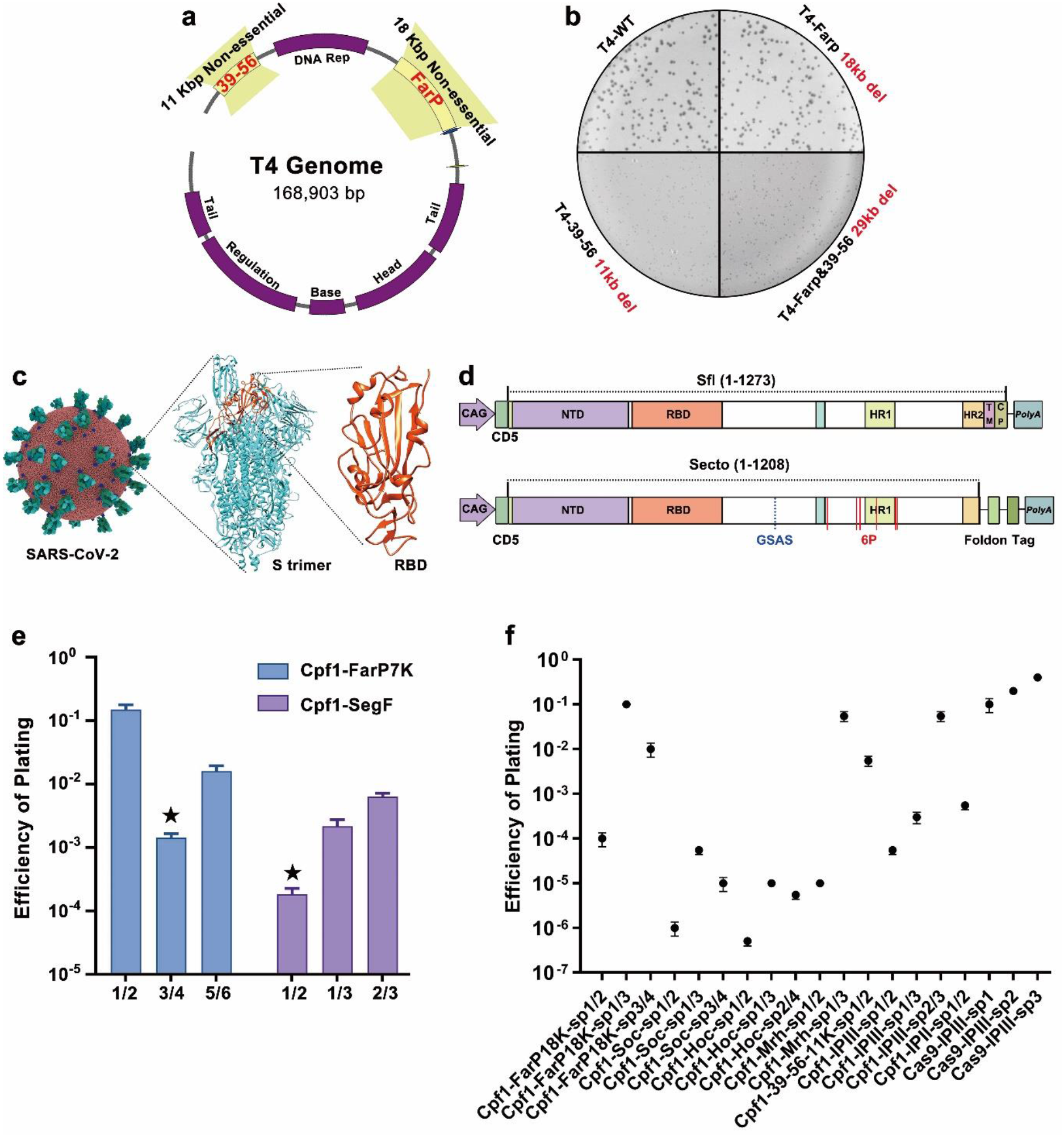
CRISPR engineering of non-essential T4 genome. **a.** Schematic showing the 18-kb nonessential segment FarP and 11-kb nonessential segment 39-56 on T4 genome. **b.** Plaque size of wild-type (WT), T4-*FarP 18 kb del.*, T4-*39-56 11 kb del.*, and T4-*FarP&39-56 29 kb del.* phages. Note the small size of T4-*39-56 11 kb del.* and T4-*FarP&39-56 29 kb del.* plaques. **c.** Structural models of SARS-CoV-2 virus, spike trimer, and receptor binding domain (RBD). **d.** Schematics of S-full length (S-fl) and S-ectodomain (S-ecto) expression cassettes used for insertion into T4 genome. **e.** Efficiency of plating of three sets of Cpf1-FarP7K spacers and three sets of Cpf1-SegF spacers. **f.** Efficiency of plating of various spacers used for T4 genome engineering in this study.

### Encapsidation of SARS-CoV-2 nucleocapsid protein (NP)

SARS-CoV-2 infected patients have been reported to generate robust NP-specific immune responses including cytotoxic T cells that might be important for protection and virus clearance^33,36.^ We incorporated NP into the T4 nanoparticle by designing a CRISPR strategy that packaged NP molecules inside phage capsid along with the genome. As NP is a nucleic acid binding protein, the packaged phage genome might provide an appropriate environment to localize this protein^37^.

During T4 phage morphogenesis, the major capsid protein gp23 assembles around a scaffolding core formed by a cluster of proteins including three nonessential, highly basic, internal proteins IPI, IPII, and IPIII. Following assembly, while most of the scaffold proteins are degraded and expelled from capsid (Fig. 3a), the IPs are cleaved only once, next to a ~10 aa N-terminal capsid targeting sequence (CTS; MKTYQEFIAE). The cleaved CTS leaves the capsid but ~1,000 molecules of IPs, or any foreign protein fused to CTS, remain in the “expanded” capsid^38^ (Fig. 3a). To replace IPs with NP, we first created an IPIII deletion phage using appropriate spacer and donor (Supplementary Fig. 2a, Fig. 3b). Then, a CTS-NP fusion sequence was transferred into this phage under the control of the native IPIII promoter (Supplementary Fig. 2a). Next, IPII was deleted to reduce protein packaging competition and increase the copy number of NP (Supplementary Fig. 2a, Fig. 3b,c). We have also introduced an amber mutation into the CTS sequence [TTT (Phe) at aa 7] because, for unknown reasons, the donor plasmid containing the WT CTS sequence was found to be toxic to *E. coli*. This CTS*am*-NP phage when grown on a tRNA amber suppressor-containing *E. coli* (*Sup*^*1*^) expressed NP and encapsidated it as demonstrated by Western blotting (WB) with NP-specific monoclonal Abs (Fig. 3c, Supplementary Fig. 2b,c). The copy number is about 70 NP molecules per phage capsid (Supplementary Fig. 2d).

**Fig. 3.**
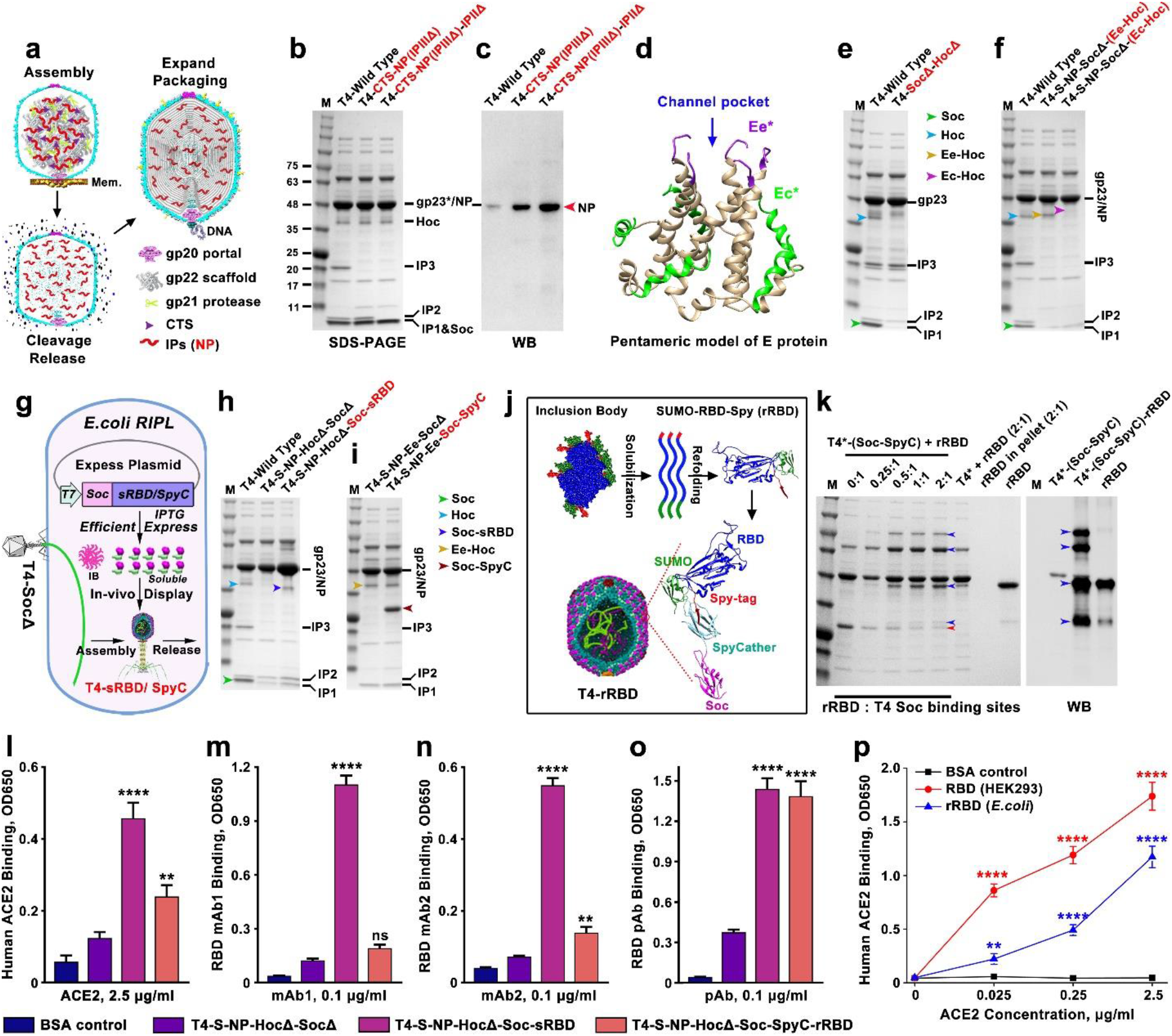
Incorporation of various SARS-CoV-2 vaccine payloads into phage T4 nanoparticle. **a.** Schematic showing steps in T4 phage head morphogenesis. Mem, *E. coli* membrane; CTS, capsid targeting sequence. **b** and **c.** SDS-PAGE and Western Blot (WB) analysis of phage particles with IPII and IPIII deletions (IPIIΔIPIIIΔ) and NP encapsidation. Since NP has a very similar molecular size to T4 major capsid protein gp23*, an NP-specific antibody was used to detect NP. **d.** Structural model of viroporin-like tetrameric assembly of CoV-2 E protein^32^. The N-terminal seven residues and C-terminal ten residues are not shown due to the lack of a corresponding segment in the structural template used for homology modeling. Ee* indicates amino acids (aa) 8-12 and Ec* indicates aa 53-65. **e.** SDS-PAGE of Hoc deletion and Soc deletion phage (HocΔSocΔ). **f.** SDS-PAGE of recombinant phages displaying Ee-Hoc or Ec-Hoc fusion proteins. **g.** Schematic showing Soc-sRBD or Soc-SpyCatcher (SpyC) *in vivo* display on T4-SocΔ capsid. Soc-sRBD or Soc-SpyCatcher expression under the control of phage T7 promoter was induced by IPTG. Most of the expressed Soc-RBD was in the inclusion body (IB). Soluble Soc-sRBD (minor amount) or Soc-SpyC can be efficiently displayed on capsid. **h.** SDS-PAGE showing ~100 copies of Soc-sRBD displayed on T4 capsid. **i.** SDS-PAGE showing ~500 copies of Soc-SpyCatcher displayed on T4 capsid. **j.** Schematic diagram showing the solubilization and refolding of SUMO (small ubiquitin like modifiers)-RBD-Spytag inclusion body. Refolded SUMO-RBD-Spytag (rRBD) protein was efficiently displayed on T4-SpyCatcher phage via Spytag-SpyCatcher bridging. **k.** Display of rRBD on the T4-SpyCacher surface at increasing ratios of rRBD molecules to capsid Soc binding sites (0:1 to 2:1). RBD specific antibody was used to verify the displayed rRBD and rRBD-SpyCatcher-Soc complexes. T4* indicates T4-S-ecto-NP-Ec-SocΔ recombinant phage. Blue and red arrows indicate rRBD/complexes and Soc-SpyCatcher, respectively. **l** to **o**. Comparison of binding of T4-sRBD, and T4-rRBD phages to soluble human ACE2 receptor (l), monoclonal antibody (mAb) 1 (human IgG Clone #bcb03, Thermo Fisher) (m), mAb2 (rabbit IgG Clone #007, Sino Bio) (n), and polyclonal antibodies (pAb) (rabbit PAb, Sino Bio) (o) using BSA and T4 phage as controls. **p.** Comparison of binding of *E. coli*-produced rRBD to human ACE2 with the HEK293-produced RBD. **P < 0.01 and ****P < 0.0001. ns, no significance, P > 0.05.

**Supplementary Fig. 2.**
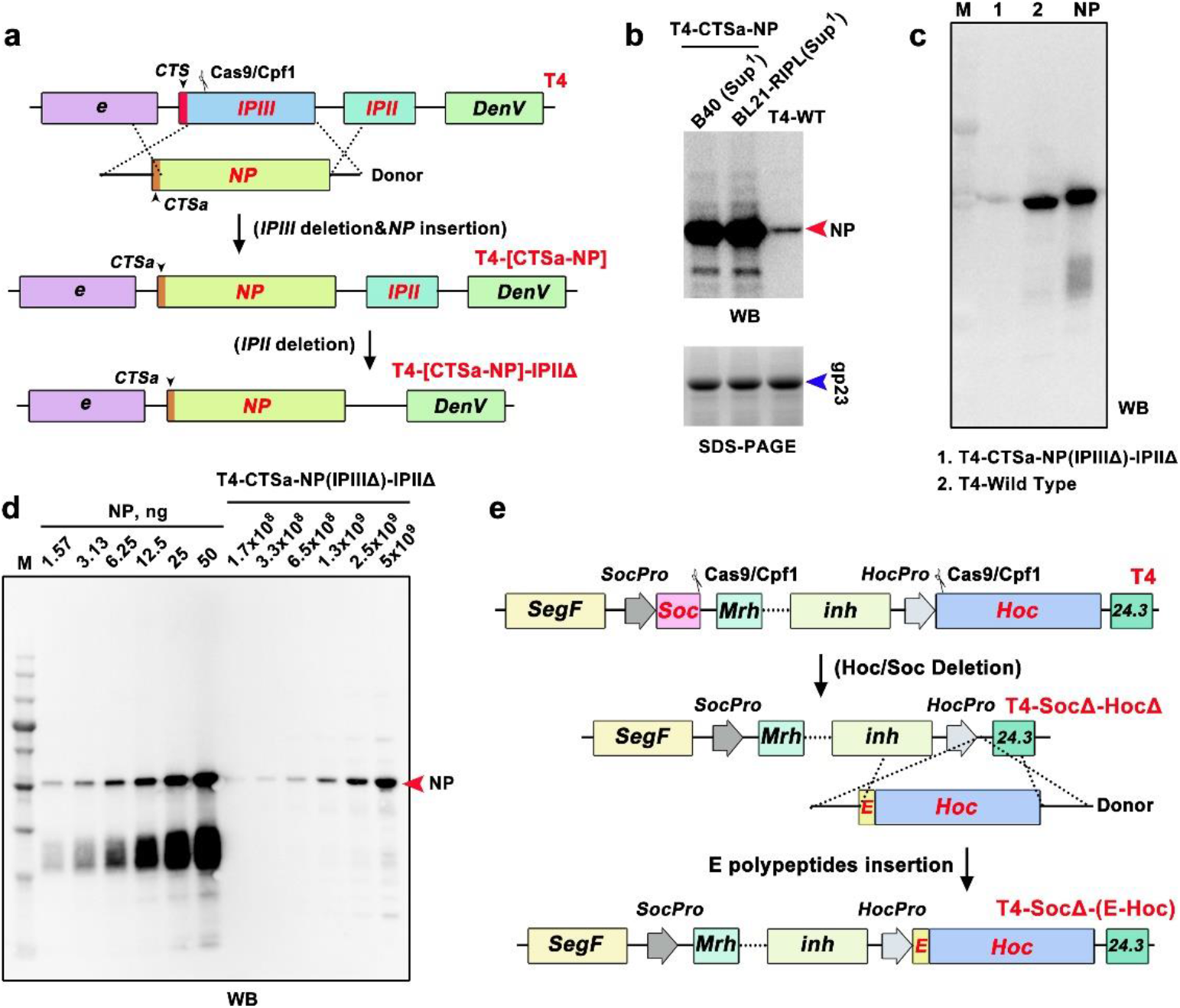
Engineering of NP encapsidation and Ee epitope display. **a.** Schematic showing the construction of T4-IPIIIΔ-IPIIΔ-CTSam-NP phage. **b.** WB showing NP expression and encapsidation in *E. coli* B40 (*Sup*^*1*^) and BL21-RIPL (*Sup*^*1*^) infected with T4-CTSa-NP phage. **c.** A second NP-specific monoclonal antibody was used to confirm the encapsidation of NP in T4-CTSa-NP phage. **d.** Quantification of the copy number of T4-encapsidated NP protein molecules by WB using commercial NP standard (Sino Bio). **e.** Schematic showing the construction of T4-SocΔ-HocΔ and T4-SocΔ-(E epitope-Hoc) phages.

### Display of SARS-CoV-2 epitopes on T4 phage

Next, we incorporated SARS-CoV-2 antigens onto the nanoparticle surface. We first deleted Hoc and Soc genes from the above recombinant phages and then inserted Hoc- and Soc-fused CoV-2 genes under the control of their respective native promoters. Upon infection, these phages would express and assemble the epitopes on T4 capsid surface (Supplementary Fig. 2e, Fig. 3d,e). We fused the gene segments corresponding to the N-terminal 12-aa “external domain” peptide (Ee) or the 18-aa peptide from the C-terminal domain (Ec) of E protein to the N-terminus of Hoc (Fig. 3d). These peptides are predicted to be exposed on the SARS-CoV-2 virion and shown to elicit T cell immune responses in humans^32^. By virtue of fusion to N-terminus of Hoc, these epitopes would be exposed at the tip of ~170 Å-long Hoc fiber^19^. The Ee and Ec recombinant phages indeed showed an upward shift of the Hoc band upon SDS-PAGE (Fig. 3f; yellow and magenta arrows). However, only the 12-aa Ee peptide was displayed at the maximum copy number, up to ~155 copies per capsid, while the 18-aa Ec peptide showed lower epitope copies and also affected phage yield (Fig. 3f).

### Display of SARS-CoV-2 receptor binding domain on T4 phage

We used a similar strategy to display CoV-2 receptor binding domain (RBD)^34^ on the capsid surface as a Soc-fusion (Supplementary Fig. 3a,b). However, the copy number of the displayed RBD was very low (Supplementary Fig. 3c). RBD contains ~82.5% non-hydrophilic residues and formed insoluble inclusion bodies in *E. coli* (Supplementary Fig. 3d,e). Numerous N- and C-terminal truncations of RBD were constructed (Supplementary Fig. 4a), however none (including the shortest 67-aa receptor binding motif^34^) had shown significant improvement in solubility and copy number (Supplementary Fig. 4b). We therefore resorted to alternative strategies to display RBD on phage capsid.

First, we constructed *E. coli* expressing Soc-RBD from a plasmid under the control of the phage T7 promoter to determine if pre-expression of Soc-RBD for a short period of time (~10 min) would keep it sufficiently soluble, which could then assemble on capsids produced during phage infection. Indeed, phage isolated from these infections showed improved display, ~100 copies of RBD (sRBD) per phage particle (Fig. 3g,h).

Second, we deployed the well-established Spytag-SpyCatcher technology^39^ to display RBD on T4 phage. The optimized SpyCatcher and Spytag from *Streptococcus pyogenes*, interact with >picomole affinity (approaching “infinite” affinity; second-order rate constant: 5.5 × 10^5^ M^−1^ s^−1^) and exquisite specificity that then leads to covalent linkage^39^. To display RBD, phage decorated with the 12.6-kDa soluble SpyCatcher was produced by growing the T4-Spike-Ee-NP-SocΔ phage on *E. coli* expressing Soc-SpyCatcher fusion protein from the T7 expression plasmid (Supplementary Fig. 3f, Fig. 3g). Phage prepared from these infections contained up to ~600 copies of Soc-SpyCatcher per capsid (Fig. 3i).

Third, RBD was expressed as SUMO-RBD-Spytag fusion protein in *E. coli*. The SUMO domain is expected to enhance the solubility of RBD^40^, but resulted in only a small improvement (Supplementary Fig. 3g). Therefore, the SUMO-RBD-Spytag protein was purified from insoluble inclusion bodies by urea denaturation and refolding (rRBD), and was displayed *in vitro* on the SpyCatcher phage (Fig. 3j). SDS-PAGE and WB of the phage particles showed that the SUMO-RBD-Spytag was efficiently captured by the SpyCatcher phage as shown by the disappearance of the SpyCatcher band and appearance of higher molecular weight band(s) (Fig. 3k). The copy number was ~300 rRBD molecules per capsid (Fig. 3k).

The sRBD and rRBD phages produced as above (T4-Spike-Ee-CTS*am*-NP-sRBD and T4-Spike-Ee-CTS*am*-NP-rRBD) bound to human ACE2 receptor protein^34^ (Fig 3l, Supplementary Fig. 5a), and to some of the RBD-specific monoclonal antibodies (mAbs) and polyclonal antibodies (pAbs) but not to all (Fig. 3m to 3o, Supplementary Fig. 5b to 5d). However, these RBDs exhibited considerably lower binding when compared to mammalian-expressed RBD suggesting that the *E. coli*-produced RBDs were not properly folded (Fig. 3p). This was also consistent with the co-purification of a 65-kDa *E. coli* GroEL chaperone with RBD indicating the presence of partially folded and/or misfolded RBD (Supplementary Fig. 3e).

**Supplementary Fig. 3.**
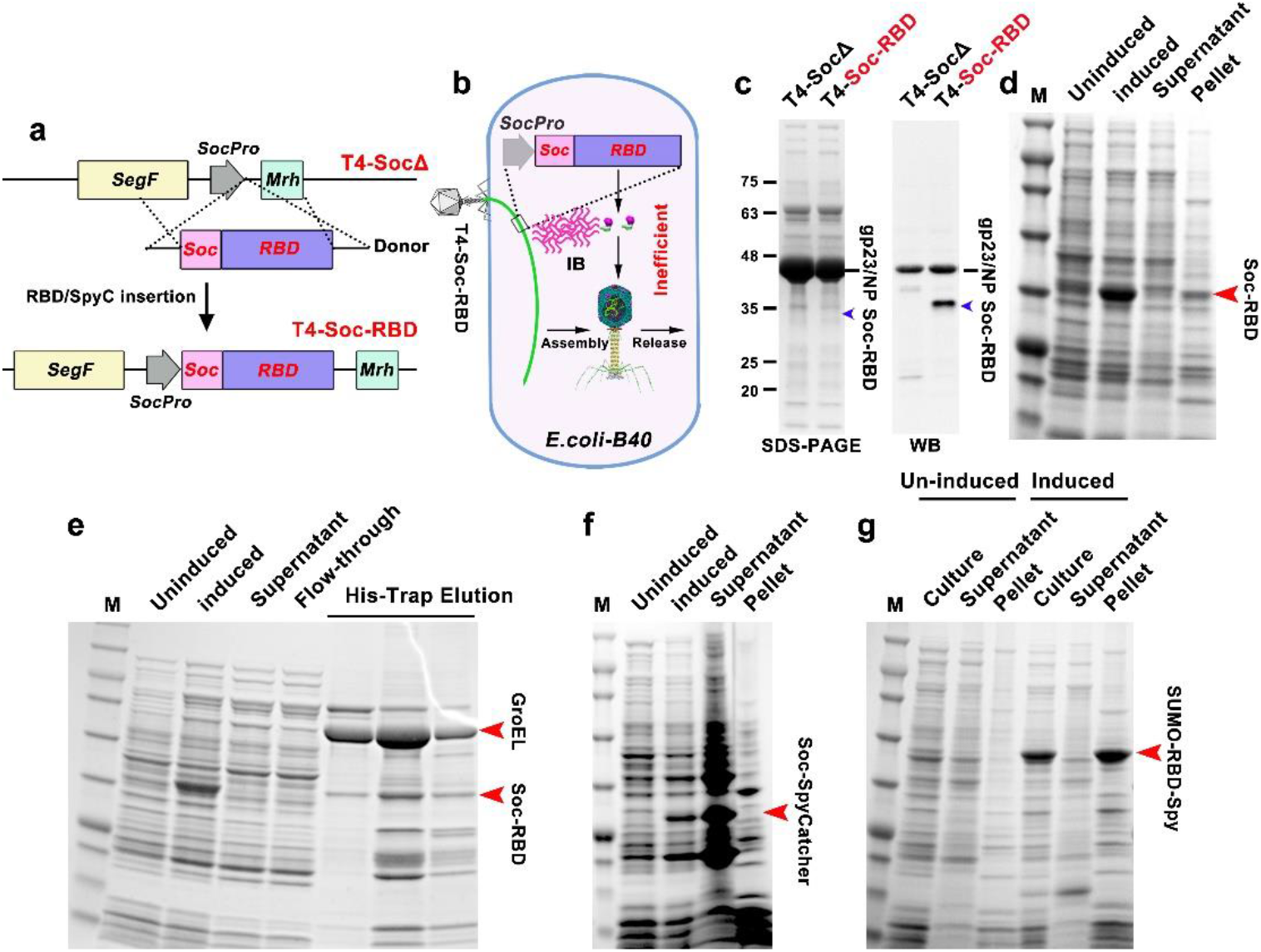
Engineering and solubility analysis of Soc-RBD, SUMO-RBD-Spytag, and Soc-SpyCatcher constructs. **a.** Schematic showing the insertion of Soc-RBD gene into phage genome at the Soc deletion site. **b and c.** Schematic (b) and SDS-PAGE/WB (c) showing inefficient *in vivo* display of *E. coli*-expressed sRBD on T4 phage. IB, inclusion body. No significant Soc-RBD band was observed by SDS-PAGE but it could be detected by WB. Anti-RBD polyclonal antibody (Sino Bio) used here also un-specifically recognized T4 gp23. **d.** Solubility analysis of Soc-RBD. The presence of Soc-RBD in the pellet and absence in the supernatant of *E. coli* lysate indicates insolubility. **e.** Very little soluble Soc-RBD was recovered after concentration of the any soluble Soc-RBD by purification on a HisTrap Ni^2+^ affinity column. No significant Soc-RBD was detected in supernatant and flow-through, but a small amount of Soc-RBD was co-eluted with *E. coli* GroEL chaperone. **f.** Solubility analysis of Soc-SpyCatcher. Soc-SpyCatcher expression was driven by the phage T7 promoter. Most of the expressed Soc-SpyCatcher protein remained in the supernatant indicating its high solubility. **g.** Solubility analysis of SUMO-RBD-Spytag. SUMO-RBD-Spytag is insoluble, similar to Soc-RBD.

**Supplementary Fig. 4.**
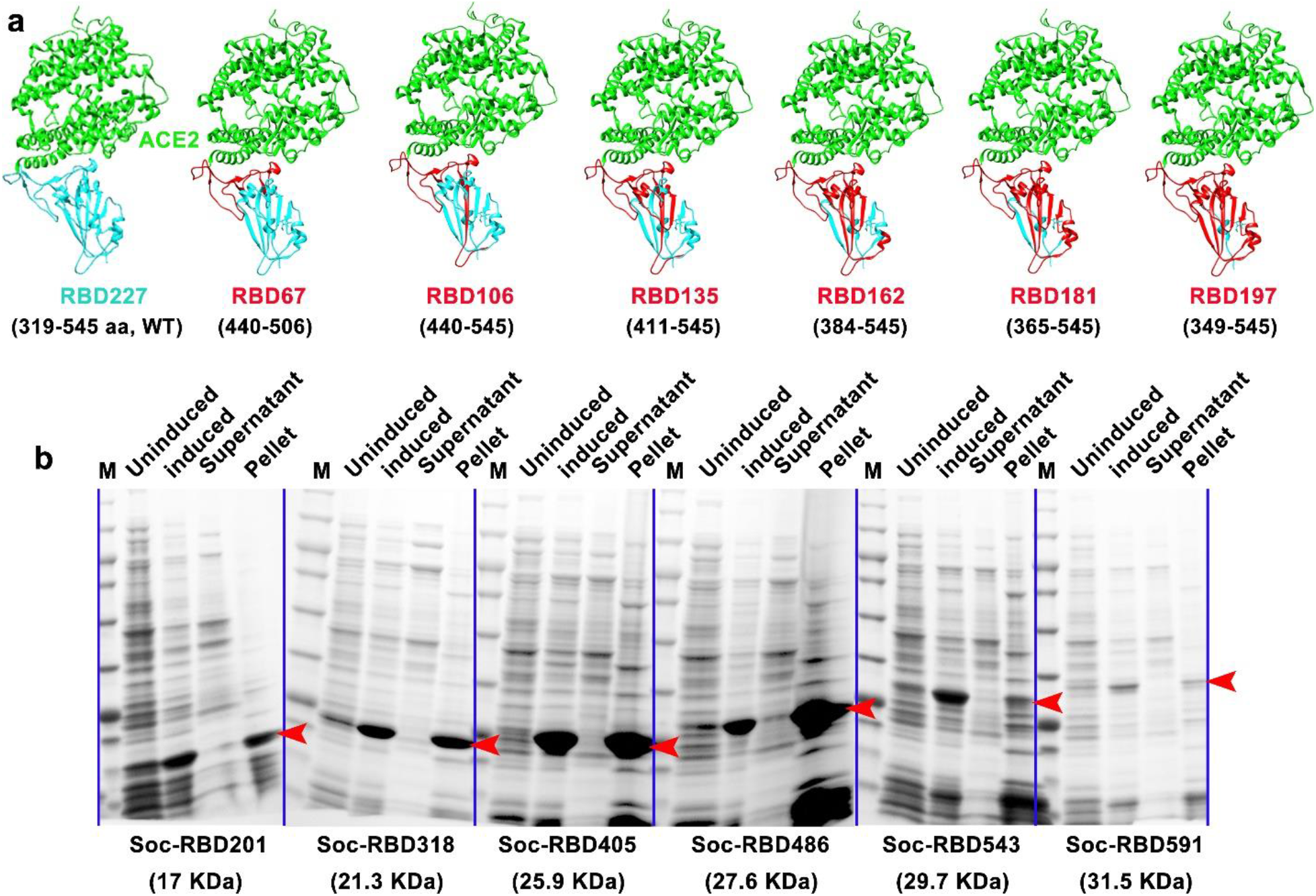
Construction and screening of various truncated SARS-CoV-2 RBDs. **a.** Structural models of recombinant WT RBD and various truncated RBDs bound to human ACE2. ACE2 is shown in green. The truncated RBD clones are shown in red and the WT RBD and deleted regions are shown in cyan. The Protein Data Bank (PDB) code for the SARS-CoV-2 RBD–ACE2 complex is 6M0J^34^. The truncated RBDs were generated using Chimera software. **b.** Solubility analysis of Soc-fused truncated RBDs after cloning and expression in *E. coli* under the control of the phage T7 promoter. After lysis of *E. coli* and centrifugation, the supernatant and pellet were analyzed by SDS-PAGE. The presence of Soc-truncated RBDs in the pellet and their absence in the supernatant demonstrated insolubility. The red arrowheads indicate the band positions of various Soc-truncated RBDs.

**Supplementary Fig. 5.**
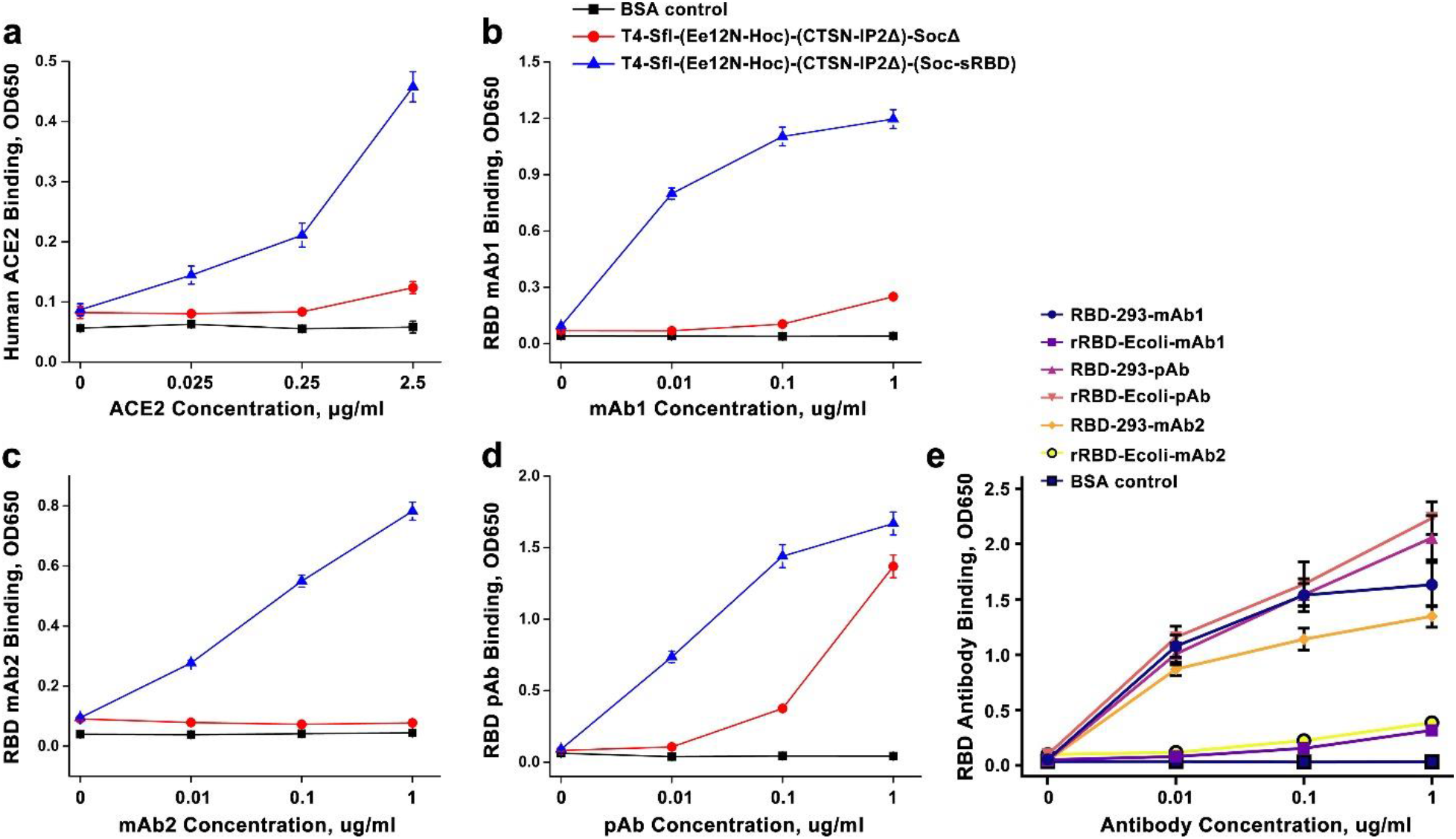
Comparison of ACE2 and RBD-antibody binding of T4-sRBD, *E.coli*-rRBD, and HEK293-RBD. **a** to **d**. ACE2 and a panel of RBD-specific antibodies used for quantification by ELISA. **e.** Comparison of *E. coli*-produced rRBD and human HEK293-produced RBD using a panel of RBD-specific mAbs and pAbs. The HEK293-RBD showed much greater binding to mAb1 and mAb2 than the *E. coli* rRBD, while binding to pAbs was similar.

### Decoration of phage T4 nanoparticles with spike ectodomain trimers

Next, we displayed spike ectodomain (S-ecto) trimers (S-trimer) (433.5 kDa)^31^ on T4-Spike-Ee-NP-SocΔ phage. The pre-fusion stabilized hexa-Pro S-ectodomain construct described as above^31^ was fused to a 16-aa spytag at the C-terminus and expressed in ExpiCHO cells. The ectodomain trimers secreted into the culture medium were purified by HisTrap affinity chromatography and size-exclusion chromatography (Supplementary Fig. 6a,b). These trimers appeared authentic and native-like because: i) they migrated predominantly as a single species and showed no nonspecific aggregation, which usually appears as a peak (or shoulder) near void volume and as a smeary high molecular weight species on native gel (Supplementary Fig. 6a,b), and ii) they bound efficiently to human ACE2 receptor (Supplementary Fig. 6c) and to conformation-specific RBD mAbs (Supplementary Fig. 5e). Importantly, the S-trimers were efficiently captured by the SpyCatcher phage produced as above. Binding was so strong that efficient assembly occurred by simple mixing of trimers and phage even at an equimolar ratio of S-trimers to T4-spycatcher molecules (Fig. 4a,b). The copy number was ~100 S-trimers per phage capsid (Fig. 4b). Cryo-EM showed decoration of T4 phage capsids with S-trimers (Fig. 4c), mimicking the orientation of spikes on SARS-CoV-2 virion^41^. The trimer-decorated T4 phage efficiently bound to human ACE2 receptor (Fig. 4d, Supplementary Fig. 7a) and when co-displayed with GFP, it decorated the ACE2-expressing HEK293 cells (Fig. 4e, Supplementary Fig. 7b, c).

**Fig. 4.**
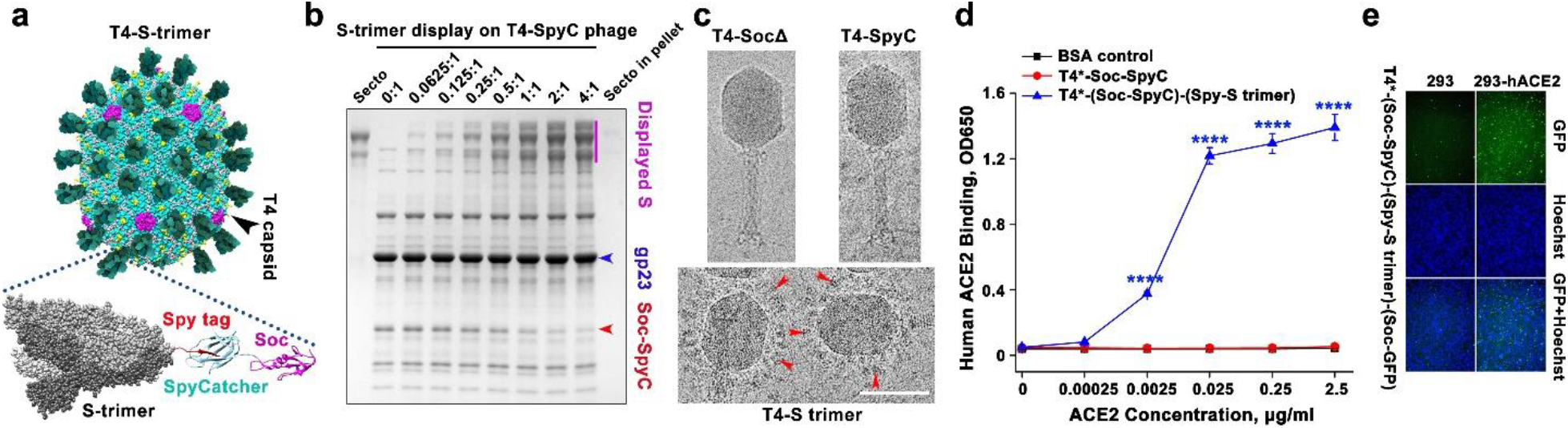
Decoration of phage T4 nanoparticles with spike ectodomain trimers. **a.** Schematic showing the decoration of phage T4 nanoparticles with spike ectodomain trimers via Spytag-SpyCatcher bridges. **b.***In vitro* assembly of S-trimers on T4-SpyCatcher phage at increasing ratios of S-trimer molecules to Soc binding sites (0:1 to 4:1). Phage and S-trimer were incubated at 4°C for 1 hr, followed by centrifugation to remove the unbound material. After two washes, the pellet was re-suspended in buffer and SDS-PAGE was performed. **c.** Representative cryo-EM images showing T4-SocΔ, T4-(Soc-SpyCatcher), and T4-(Soc-SpyCatcher)-S-trimers phages. The red arrowhead indicates the representative S-trimer displayed on phage. Bar = 100 nm. **d.** ELISA analysis of T4-S-trimer phage binding to ACE2 at various ACE2 concentrations. ****P < 0.0001. **e.** Binding of T4-S-trimer-GFP phage to HEK293 cells expressing human ACE2. The nucleus was stained with Hoechst. T4* indicates T4-(S-ecto)-RBD-NP-Ee-SocΔ phage used for display (S-ecto and RBD refer sot insertions of gene expression cassettes).

**Supplementary Fig. 6.**
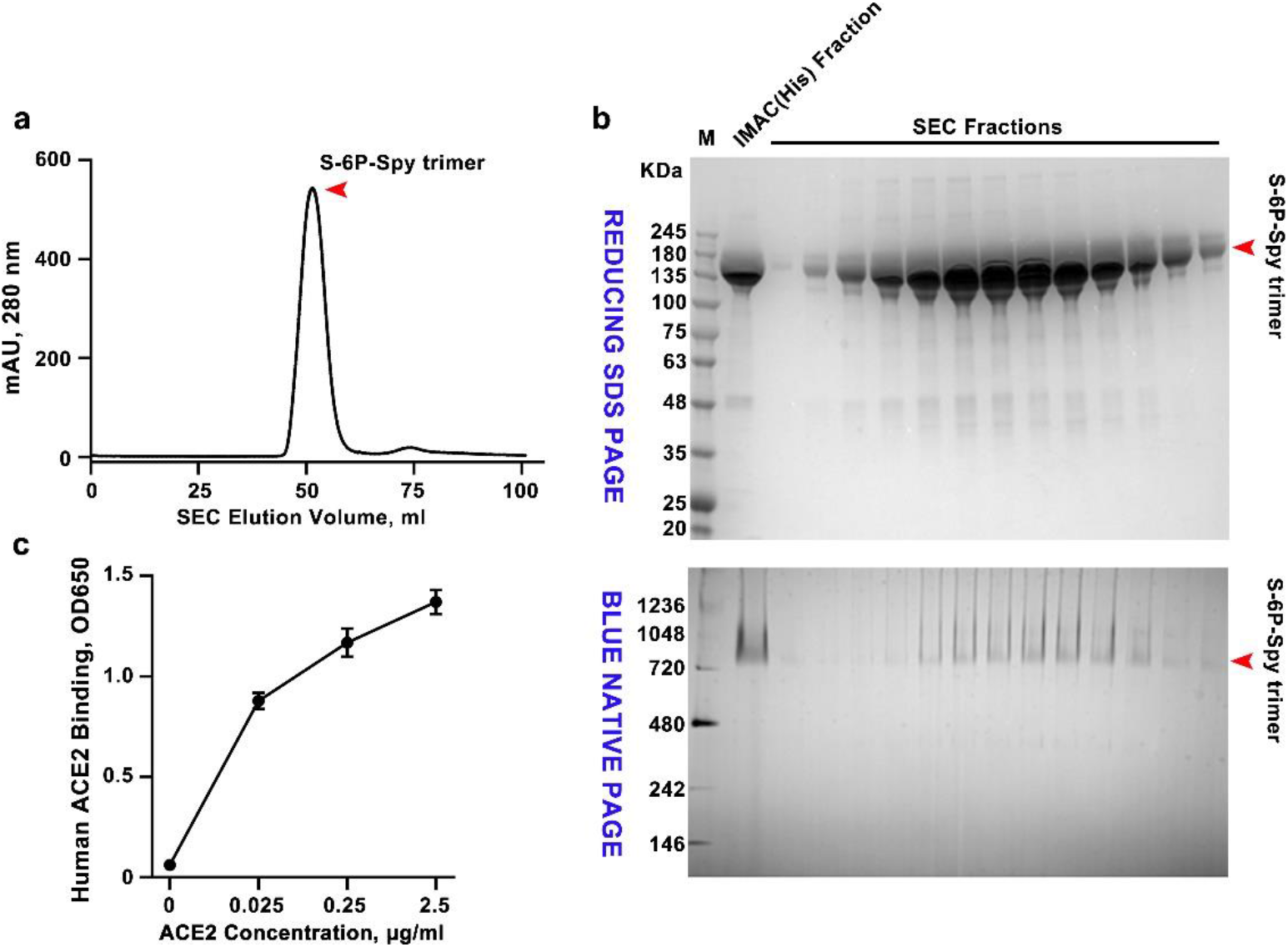
Purification and characterization of S-ecto-spytag trimers from ExpiCHO cells. **a.** Size-exclusion chromatography (SEC) elution profile of S-ecto-spytag (S-ecto-spy) trimers. HisTrap affinity purified S-ecto-spy protein from 250 ml of transfected ExpiCHO cells was loaded on Superdex 200 prep-grade SEC column. S-trimers yield was ~50 mg per 1 L culture. **b**. Reducing SDS-PAGE (top) and BLUE NATIVE-PAGE (bottom) patterns of SEC-purified trimer fractions. The molecular weight standards (M) in kDa are shown on the left of the gels. IMAC (Immobilized Metal Affinity Chromatography, His) fraction is the material from affinity purification of culture supernatant on a HisTrap column, which was then loaded on the SEC column. **c.** ELISA analysis showing binding of purified S-trimers to human ACE2 at various ACE2 concentrations.

**Supplementary Fig. 7.**
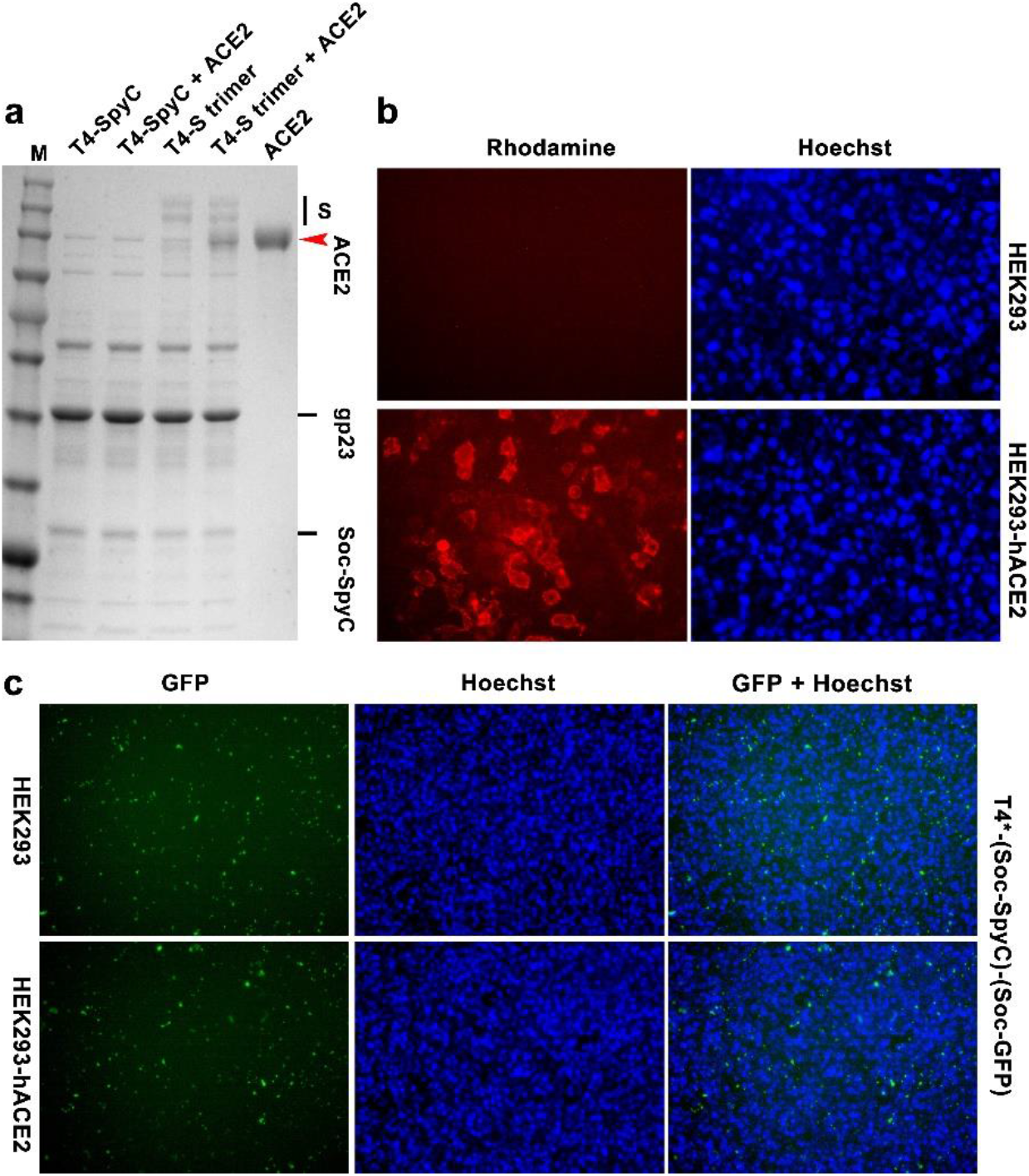
Binding of T4 phage-decorated S-trimers to ACE2-expressing HEK 293 cells. **a.** Co-sedimentation assay showing the capture of ACE2 by T4 phage-decorated S trimers. T4-S-trimer particles and ACE2 were incubated at equimolar ratio for 1 hr at 4°C, followed by high speed centrifugation. After two washes, the pellet was re-suspended in buffer and SDS-PAGE was performed. Presence of ACE2 in the pellet was found with these phage particles but not with the control phage lacking S-trimers. **b.** Immunofluorescence assay showing expression of ACE2 on 293 cells. Two days after ACE2 plasmid transfection, HEK293 cells were incubated with RBD, followed by anti-RBD antibody and Rhodamine-conjugated second antibody. **c.** Lack of binding of T4-GFP control phage (without S-trimers) to ACE2-293 cells. No difference in fluorescence was observed. The nuclei were stained with Hoechst. T4* indicates T4-(S-ecto)-RBD-NP-Ee-SocΔ.

### Immunogenicity and protective efficacy of T4-SARS-CoV-2 vaccine candidates

The T4-CoV-2 vaccine candidates generated as above by sequential engineering (Supplementary Fig. 8a,b) were screened for their immunogenicity and protective efficacy in a mouse model. BALB/c mice were immunized at weeks 0, 3, and 6 with CsCl-purified phage particles (Fig. 5a,b) and sera were analyzed by a series of immunological assays.

**Fig. 5.**
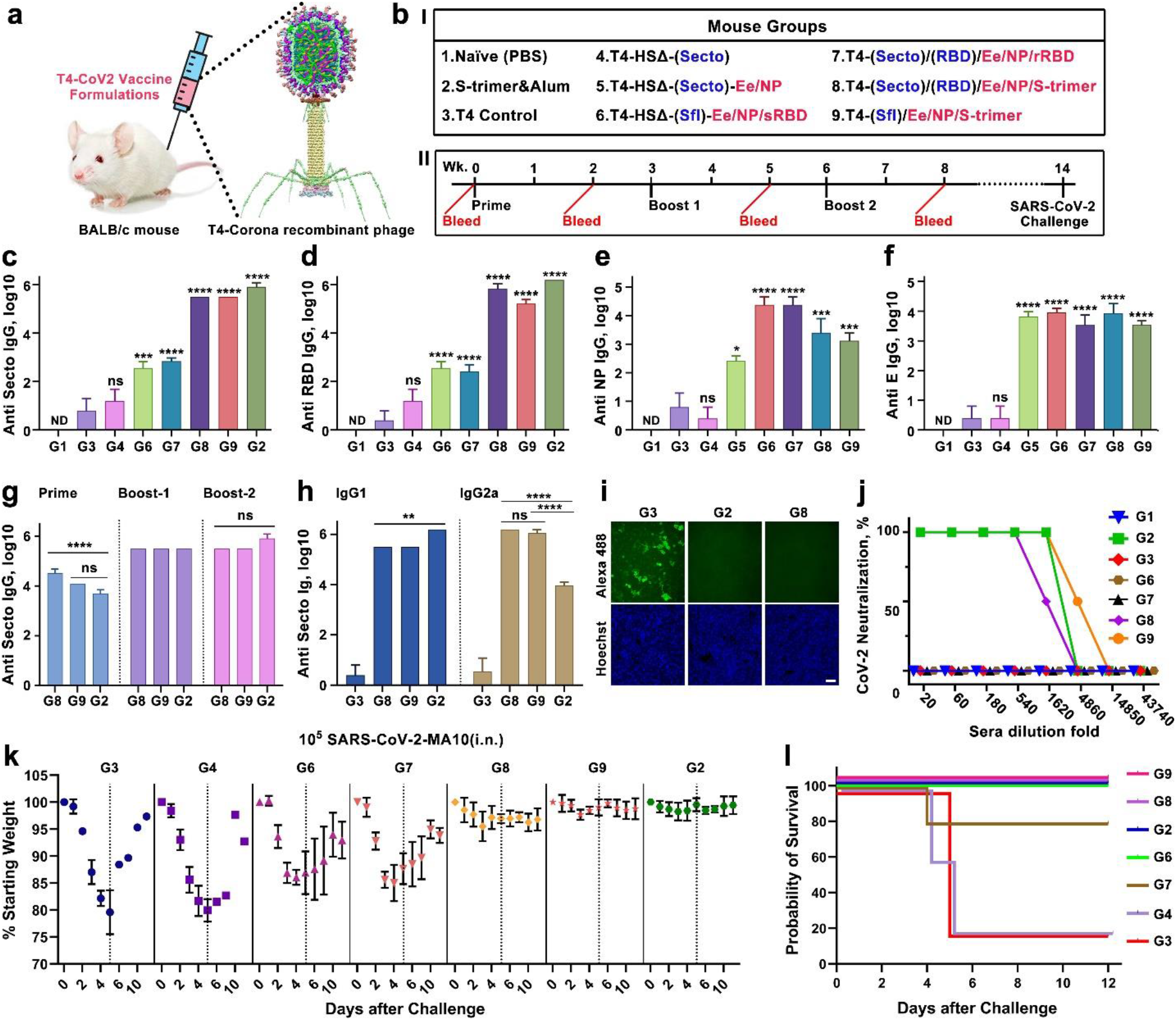
Immunogenicity and protective efficacy of T4-SARS-CoV-2 vaccine candidates in mice. **a.** Schematic diagram showing BALB/c mice immunized by the intramuscular (i.m.) route using T4-SARS-CoV-2 vaccine formulations. **b. I.** Formulations and mouse groups used for vaccinations. HSΔ indicates Hoc deletion and Soc deletion. Blue color (S-ecto, S-fl, and RBD) indicates the insertion of mammalian gene expression cassette into T4 genome as DNA vaccine. Red color indicates the capsid-displayed Ee, S-trimers, or *E.coli*-produced rRBD or sRBD protein, or the capsid-encapsidated NP protein. Naïve mice and mice immunized with the phage lacking any CoV-2 genes served as negative controls whereas mice immunized with S-trimers adjuvanted with Alhydrogel served as a positive control. **II.** Prime-boost immunization scheme. BALB/c mice (5 per group) were immunized on weeks 0, 3, and 6 and challenged intranasally (i.n.) with a mouse-adapted SARS-CoV-2 strain (SARS-CoV-2 MA10)^47^ on week 14. **c** to **f**. The boost-2 sera (week 8 bleeding) from various groups were assessed by ELISA for antigen-specific IgG antibody titers (endpoint) against S-ecto (c), RBD (d), NP (e), and E (f). *P < 0.05, **P < 0.01, ***P < 0.001, and ****P < 0.0001, compared with phage control group G3. ns, no significance, P > 0.05. ND, not detected. **g.** Measurement of anti-S-ecto IgG antibody titers in sera from S-trimers-Alhydrogel (G2) group and T4-S-trimers (G8 and G9) groups at weeks 2 (prime), 5 (boost-1), and 8 (boost-2). ***P < 0.001. **h.** Comparison of anti-S-ecto IgG1 and IgG2a subtype antibody titers in sera from S-trimers-Alhydrogel (G2) group and T4-S-trimers (G8 and G9) groups at 8 weeks (boost-2). **P < 0.01 and ****P < 0.0001. **i.** Blocking of native RBD protein binding to HEK293-ACE2 by sera from phage control group (G3), S-trimers-Alhydrogel group (G2), and T4-S-trimers group (G8). The sera were diluted 500-fold. RBD binding to HEK293-ACE2 was detected by Alexa 488 conjugated secondary antibody (primary antibody: anti-RBD human IgG). **j.** Neutralization antibody measurement. Infection of Vero E6 cells by SARS-CoV-2 live virus was determined in the presence of mouse sera at a series of threefold dilutions starting from 1:20. **k.** Percentage starting body weight of immunized mice at days post infection with 10^5^ PFU SARS-CoV-2 MA10 (i.n.). Dotted line represents percentage starting weight at day 5 post infection, in which mice showed maximum weight loss in control groups. In groups G3 and G4, only 20% of mice survived after day 5. Thus, data presented after day 5 are biased toward minor survivors. The data were presented as means ± SD. **l.** Survival rate of mice against SARS-CoV-2 MA10 challenge.

SARS-CoV-2-specific antibody titers were determined by ELISA using purified proteins (S-ecto, RBD, NP, or E) as coating antigens. Recombinant phages that delivered CoV-2 DNA alone did not induce significant titers of spike protein-specific antibodies (Fig. 5c,d, Supplementary Fig. 9a to 9d). However, when the same were boosted once with phage nanoparticles displayed with S-trimers, significant antibody titers were elicited (Supplementary Fig. 10a).

Without any adjuvant, the T4 nanoparticles stimulated strong antibody titers to phage-delivered proteins/peptides, either displayed on surface or packaged inside. These include RBD- and spike-specific antibodies, E-specific antibodies, and NP-specific antibodies (Fig. 5c to 5f). However, the highest titers, up to an endpoint titer of ~1.5 × 10^6^, were obtained with phage nanoparticles decorated with S-trimers (Fig. 5c to 5f). Most of these antibodies are RBD-specific since no significant difference was observed between the endpoint titers obtained by using either RBD or S-trimers as the coating antigen (Fig 5c,d). This result is consistent with recent reports that the immunodominant RBD comprises multiple distinct antigenic sites and is the target of most neutralizing activity in COVID-19 convalescent sera^42, 43^.

Furthermore, the antibodies elicited were conformation-specific. For instance, the antibodies elicited against sRBD or rRBD displayed on phage reacted poorly with the mammalian-expressed S-trimer or RBD (Fig 5c,d), consistent with the antigenicity data described above that these also reacted poorly with ACE2 and conformation-specific mAbs (Fig. 3p, Supplementary Fig. 5e). Similarly, the antibodies elicited against T4-dispalyed S-trimers reacted poorly with the *E.coli*-produced RBD (end point titer of ~10^2^ using *E. coli* RBD as the coating antigen vs ~10^5^ using mammalian RBD) (Supplementary Fig. 10d).

The spike-specific titers elicited by phage-decorated trimers without any adjuvant were as high as those generated with Alhydrogel adjuvant (Fig. 5c,d,g). Furthermore, notably, IgG subclass data indicated that phage nanoparticles stimulated both humoral (T_H_2) and cellular (T_H_1) arms of the immune system. In mice, IgG2a subclass represents T_H_1 response whereas IgG1 class reflects T_H_2 response. The adjuvant-free T4 nanoparticles mounted high levels of both IgG1 and IgG2a classes against all three SARS-CoV-2 antigens; spike/RBD, E, and NP, whereas the Alhydrogel-adjuvanted trimers predominantly elicited T_H_2-derived IgG1 class antibodies (Fig. 5h, Supplementary Fig. 9a to 9j). In fact, the T_H_1-derived IgG2a antibodies were higher than the T_H_2 derived IgG1 antibodies in T4 groups, whereas the Alhydrogel-adjuvanted mice induced ≥2 orders of magnitude lower IgG2a antibodies (Fig. 5h, Supplementary Fig. 9i,j). While T_H_2-biased responses may lead to lung injury via eosinophilic infiltrates, the T_H_1-type responses are proposed to alleviate potential lung immunopathology and reduce the potential for disease enhancement^44, 45^. The T4 nanoparticle vaccine with a balanced T_H_1 and T_H_2 responses might therefore be optimal for safety and virus clearance. This point however requires further investigation.

The T4-stimulated spike-specific antibodies blocked binding of RBD to human ACE-2 expressed on HEK293 cells in a dose-dependent manner (Fig. 5i, Supplementary Fig. 9k). Importantly, these antibodies also exhibited strong virus neutralizing activity as determined by Vero E6 cell cytopathic assay using the live BSL-3 SARS-CoV-2 US-WA-1/2020 strain, the first patient isolate obtained through the CDC^46^ (Fig. 5j). The neutralization titers correlated well with protective efficacy when mice were challenged with mouse-adapted BSL-3 SARS-CoV-2 MA10 virus^47^ (Fig. 5k,l).

Upon challenge, the naïve and T4 control mice showed rapid decline in weight, up to 25% of their starting weight in five days due to acute viral infection, and were moribund (humanely euthanized) or succumbed to infection. After day 5, the surviving mice began to re-gain weight and recovery from infection during the next several days (Fig. 5k,l). This rapid weight loss resulted in ~80% mortality rate in the control animals (Fig. 5k,l). None of the groups receiving the spike DNA vaccine alone and/or CoV-2 antigens other than spike trimers showed significant protection, closely correlating to the lack or extremely weak neutralization antibody responses. However, *E. coli* RBD groups (G6 and G7) showed less weight loss and higher survival rate than the susceptible control groups (G3 and G4) (Fig. 5k,l). On the other hand, mice vaccinated with T4-decorated trimers exhibited profound neutralization antibody responses and were fully protected from the acute onset of morbidity (weight loss and other signs of illness) and mortality. The weight loss, if any, was insignificant for both these groups, as well as the positive control group vaccinated with Alhydrogel-adjuvanted trimers (Fig. 5k,l). Additionally, those mice boosted with one dose of T4-trimers also showed partial protection. The weight loss was in between unprotected and protected groups, with a milder weight loss than the unimmunized and challenged controls, clearly correlating protection with spike-specific antibodies (Supplementary Fig. 10b,c).

Next, we evaluated the most effective T4 phage-decorated S-trimers vaccine in a second animal model, the New Zealand White rabbit (Fig. 6a). This vaccine, again without any adjuvant, by just two immunizations at days 0 and 15, elicited robust spike- and RBD-specific antibodies and virus neutralization titers in rabbits that are ~5-6 times greater than those obtained in mice, (Fig. 6b to 6d, Supplementary Fig. 11). Likewise, inclusion of displayed Ee peptide and packaged NP into the nanoparticle broadened the immune responses (Fig. 6e,f) while addition of a T_H_1-biased adjuvant Alhydroxyquim-II only slightly enhanced the antibody titers (Fig. 6b to 6d).

**Fig. 6.**
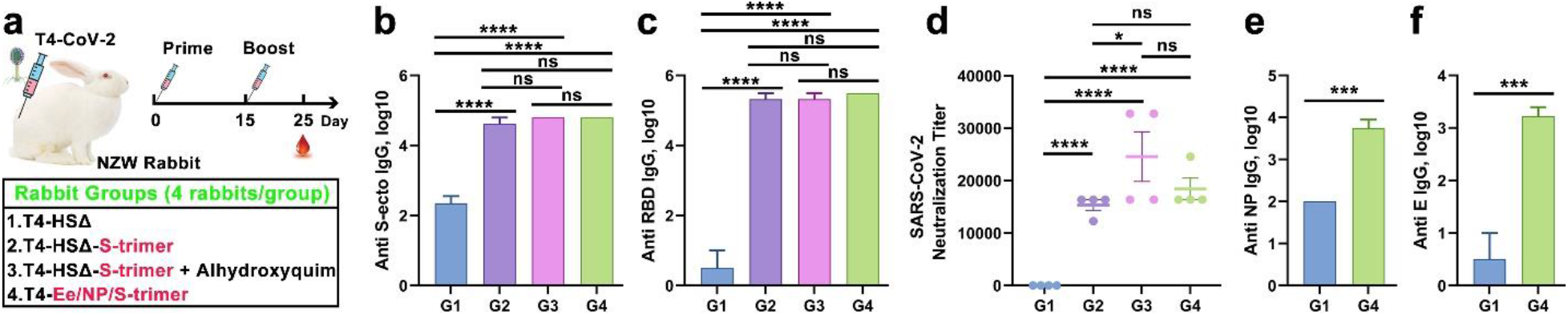
Immunogenicity and virus neutralization responses of the T4-S-trimers vaccine in New Zealand White rabbit model. **a.** Schematic diagram showing formulations, groups, and prime-single boost immunization scheme for intramuscular vaccinations of New Zealand White (NZW) rabbits. HSΔ indicates Hoc deletion and Soc deletion recombinant phage. Red color indicates the capsid-displayed Ee, S-trimers, or the capsid-encapsidated NP protein. Rabbits immunized with HSΔ phage served as negative control. **b** and **c**. boost sera (10 days after boost) were assessed by ELISA for antigen-specific IgG antibody titers against S-ecto (b) and RBD (c). ****P < 0.0001. ns, no significance, P > 0.05. **d.** Serial dilutions of serum from immunized rabbits were assessed for neutralization of live BSL-3 strain SARS-CoV-2 US-WA-1/2020. Neutralization titers were calculated as the reciprocal dilution where infection (cytopathic effect) was reduced by more than 90% relative to infection in the absence of serum. **e** and **f**. Comparison of G1 (control phage) and G4 (Ee-NP-S trimer displayed phage) for anti-NP (e) and anti-E (f) IgG antibody titers. ***P < 0.001.

Together, these datasets allowed selection of a highly effective T4 phage vaccine candidate that generated robust virus neutralization titers in two different animal models, mouse and rabbit, and conferred complete protection against acute viral infection in mice.

**Supplementary Fig. 8.**
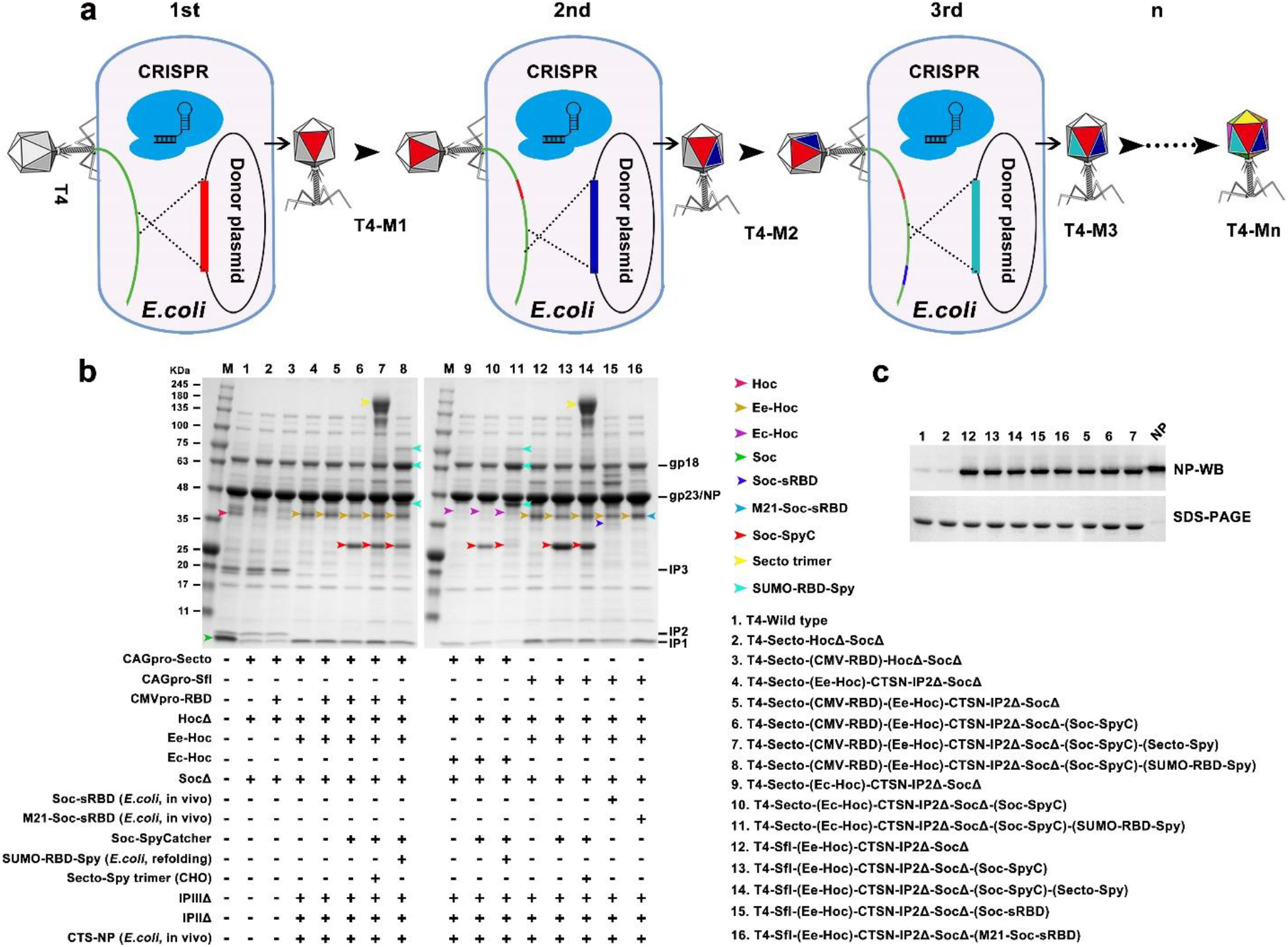
A pipeline of SARS-CoV-2 vaccine candidates generated by sequential CRISPR engineering. **a.** Schematic showing a representative sequence in which the WT phage was used as a starting infection of CRISPR *E. coli* containing spacer 1 and donor 1. The resultant T4-mutant 1 (T4-M1) was used to infect bacteria containing spacer 2 and donor 2 to produce recombinant T4-mutant 2 (T4-M2) which has two insertion/deletion mutations, and so forth. By sequential CRISPR engineering and simple phage infections, recombinant phages with multiple desired mutations were created. Each color on phage capsid here represents a mutation. **b.** One example of sequential phage CRISPR engineering for creating the T4-SARS-CoV-2 nanovaccine. Numerous CoV-2 components, including CAGpromoter-S-ecto insertion, CAGpromoter-S-fl insertion, CMVpromoter-RBD insertion, Hoc deletion, Ee-Hoc insertion, Ec-Hoc insertion, Soc deletion, Soc-sRBD display, M21-Soc-sRBD display, Soc-SpyCatcher display, refolding SUMO-RBD-Spy display, S-trimer display, IPIII deletion, IPII deletion, and NP encapsidation, were permutated and combined as needed. The resultant SARS-CoV-2 vaccine candidates were characterized by PCR, DNA sequencing and/or SDS-PAGE, and some of these were then tested in a mouse study. M21 indicates a potential T cell 21 aa epitope (SYFIASFRLFARTRSMWSFNP) from SARS-CoV-2 membrane protein. **c.** WB showing NP protein encapsidation in the phages containing CTSam-NP insertion at IPIII deletion site.

**Supplementary Fig. 9.**
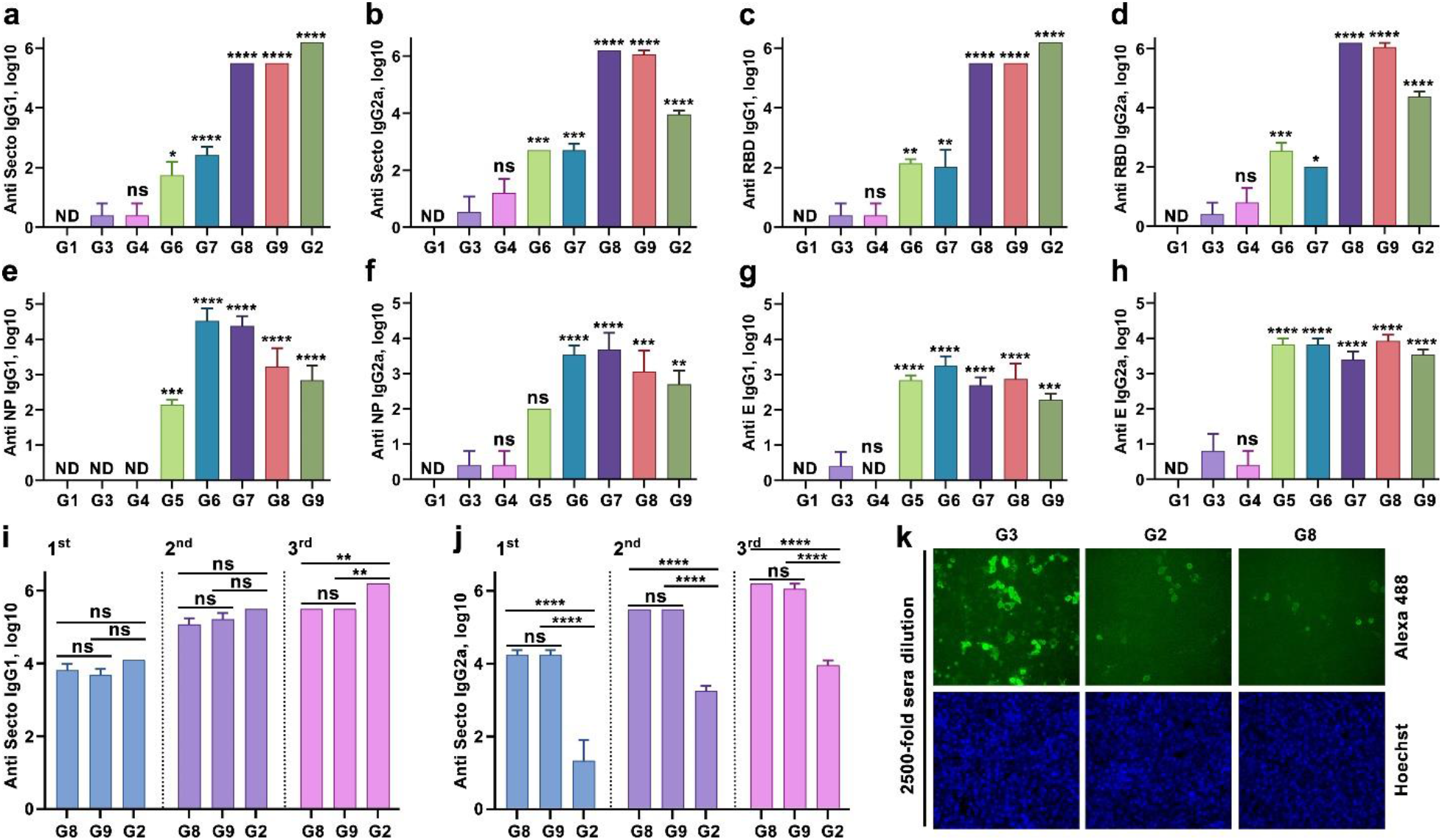
Serum antibody responses in various T4-SARS-CoV-2 immunized mice. **a** and **b.** Anti-S-ecto IgG1 (a) and IgG2a (b) antibody titers in the boost-2 sera (week 8 bleeding) from various groups. **c** and **d.** Anti-RBD IgG1 (c) and IgG2a (d) antibody titers in the boost-2 sera. **e** and **f**. Anti-NP IgG1 (e) and IgG2a (f) antibody titers in the boost-2 sera. **g** and **h**. Anti-E IgG1 (g) and IgG2a (h) antibody titers in the boost-2 sera. *P < 0.05, **P < 0.01, ***P < 0.001, and ****P < 0.0001, compared with phage control group G3. ns, no significance, P > 0.05. ND, not detected. **i** and **j.** Anti-S-ecto IgG1 (i) and IgG2a (j) antibody titers in the sera from S-trimer & Alhydrogel (G2) group and T4-S-trimer (G8 and G9) groups at 2 weeks (prime), 5 weeks (boost-1), and 8 weeks (boost-2). **P < 0.01 and ****P < 0.01. **k.** Blocking of native RBD protein binding to HEK293-ACE2 by 2500-fold diluted sera. Sera from phage control group (G3), S-trimers-Alhydrogel group (G2), and T4-S-trimers group (G8) were compared. The RBD binding to ACE2 was detected by Alexa 488 conjugated secondary antibody.

**Supplementary Fig. 10.**
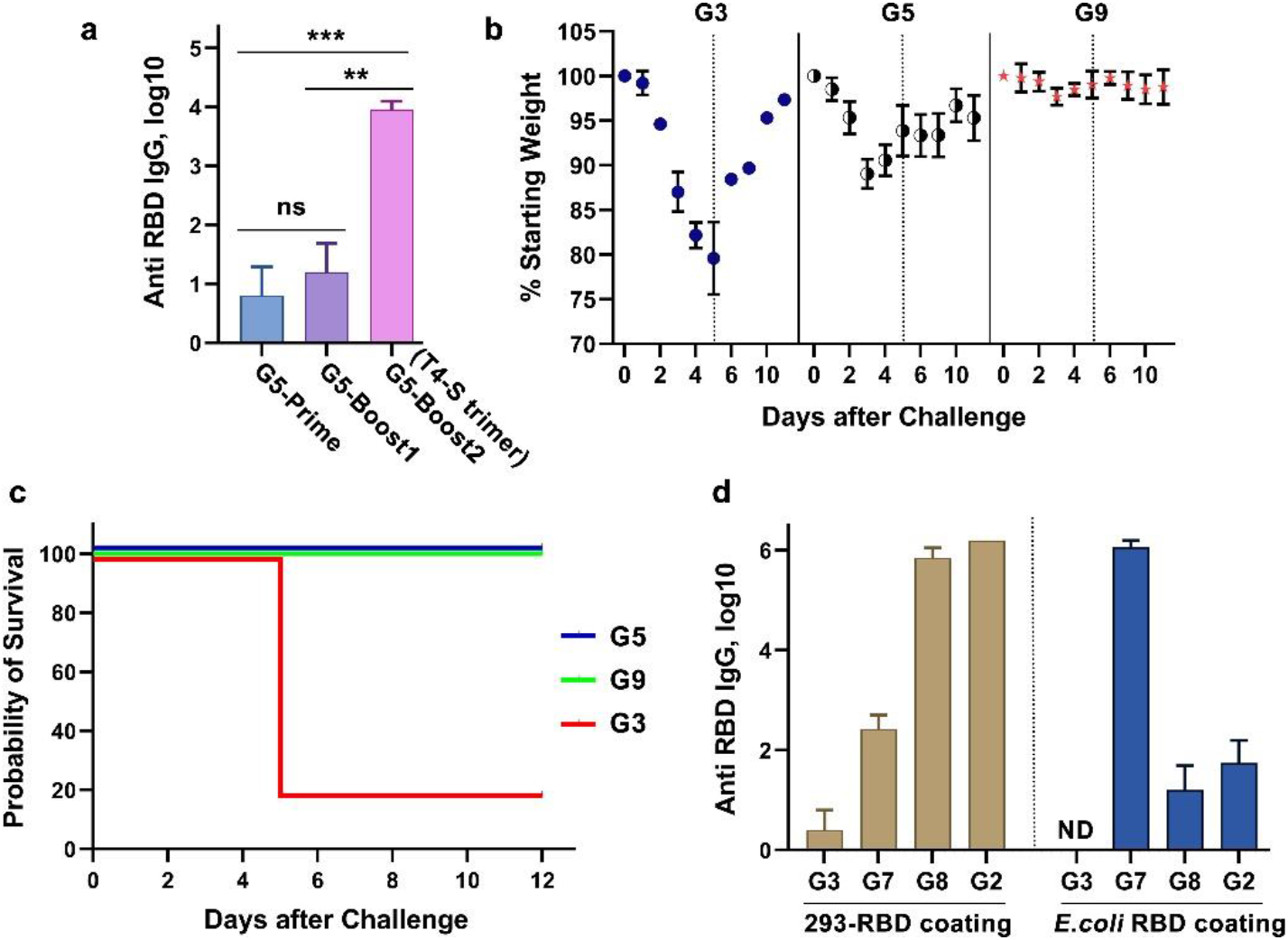
Immune responses of T4-SARS-CoV-2 immunized mice. **a.** Anti-RBD IgG antibody titers in the sera from group G5 (T4-HocΔ-SocΔ-S-ecto-Ee-NP) at weeks 2 (prime), 5 (boost-1), and 8 (boost-2). For boost-2, T4-S-trimers particles were used. **P < 0.01 and ***P < 0.001. **b.** Body weight of immunized mice from groups G3 (phage control), G5 (T4-S DNA plus T4-S-trimers boost-2), and G9 (T4-S-trimers) at days post-challenge with 10^5^ pfu of SARS-CoV-2 MA10 virus (intranasal inoculation). **c.** Survival rate of groups G3, G5, and G9 after virus challenge. **d.** Comparison of anti-RBD IgG antibody titers by ELISA using HEK293-produced RBD or *E. coli*-produced RBD as coating antigens in groups G3 (phage control), G7 (rRBD displayed T4), G8 (T4-S-trimers), and G2 (S-trimers-Alhydrogel).

**Supplementary Fig. 11.**
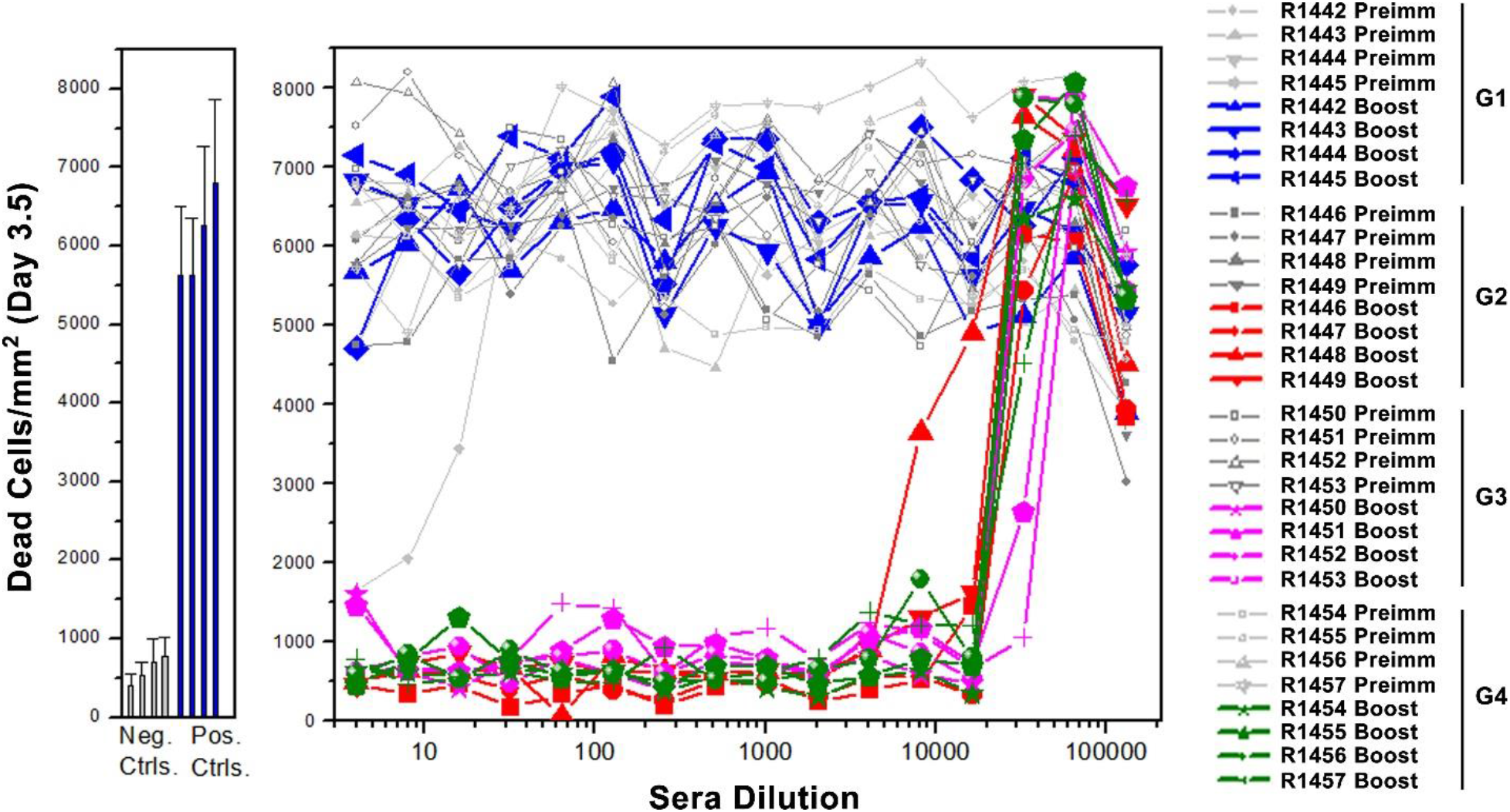
Virus neutralization titers of rabbit sera. Infection of Vero E6 cells by live SARS-CoV-2 US-WA-1/2020 was determined in the presence of rabbit sera at a series of two-fold dilutions starting from 1:4. Culture medium only and CoV-2 virus only were used as negative and positive controls, respectively. R1442 to R1457 refer to tag numbers of rabbits. The data in control groups were presented as means ± SD of 32 wells. The data in rabbit sera groups were shown as means of duplicates.

## Discussion

We have developed a “universal” vaccine platform centered around bacteriophage T4 nanoparticle and CRISPR engineering (Figure 1; Video). A number of special features demonstrated here would make this a powerful platform to rapidly generate vaccine candidates against any emerging and pandemic pathogen in the future.

First, a series of recombinant phages containing SARS-CoV-2 gene insertions were created in days, by CRISPR genome engineering using a combination of type II Cas9 and type V Cpf1 nucleases. This combination provided built-in choices for spacers as well as for efficient cleavage of T4 genome that is extensively modified by cytosine hydroxymethylation and glycosylation, to attain near 100% success^21, 22^.

Second, a large amount of genetic and structural space available in phage T4^35^ was exploited to incorporate payloads containing CoV-2 DNAs, peptides, proteins, and/or complexes into the same nanoparticle. For instance, we have inserted ~6.5 kb full-length spike gene expression cassette, 2.7 kb RBD gene expression cassette, and 1.3 kb nucleocapsid gene into the same genome by replacing nonessential genetic material of T4. It was further expanded by inserting Hoc and Soc fusions, and/or replacing additional nonessential segments that span across the phage genome. In addition, up to 155 copies of a 12-aa Ee peptide and ~100 copies of 433-kDa S-trimers were displayed on the same nanoparticle while ~70 molecules of 50-kDa NP were co-packaged with genome core. These represent unprecedented engineering capability and payload complexity by a vaccine delivery vehicle.

Third, different compartments of phage nanostructure were utilized for placing different vaccine cargos. The tips of Hoc fibers with ~170Å reach were used to display short 12-aa Ee peptide epitopes as this would allow efficient capture by antigen presenting cells, and B and T cells^29^. Indeed, these elicited strong antibody titers. At the same time, NP was co-packaged with genome as this mimics NP’s natural nucleic acid environment^2, 37^.

Fourth, the T4 platform was readily adapted to mammalian-expressed proteins through highly efficient SpyCatcher-SpyTag bridging^39^. This might be essential for proper folding and glycosylation of some pathogen antigens, as in the case of spike trimers^30^. As the Cryo-EM images showed, such spikes anchored to capsid mimic the spikes on SARS-CoV-2 virion^41^ providing a native-like context for stimulating effective immune responses.

Fifth, alternative strategies such as pre-expression from a plasmid and display through subsequent phage infection provided additional advantages to enhance copy number and better control over folding. As demonstrated, this approach yielded 3-6-fold higher copy number of Soc-SpyCatcher on phage capsid. However, the hydrophobic RBD remained poorly folded.

Sixth, our sequential engineering generated a pipeline of SARS-CoV-2 vaccine candidates in mere weeks, and allowed down-selection of the best vaccine candidate, the phage-decorated trimers, in a single mouse experiment. However, why the DNA-alone vaccines failed to elicit immune responses remains unclear and requires further investigation.

Seventh, unlike the DNAs, all the phage-delivered peptides/proteins, whether displayed on surface or packaged inside, generated robust immune responses. The strong virus neutralization titers and ACE-2 blocking titers elicited by T4-delivered trimers in two different animal models, mouse and rabbit, are particularly noteworthy. These responses correlated with protective efficacy where the vaccinated mice were completely protected from acute viral infection.

Eighth, T4 nanoparticle vaccines generated balanced T_H_1 and T_H_2 derived antibody responses against all three CoV-2 antigens tested. In fact, T4 seems to have a slight T_H_1-bias. These are desirable properties for protection and distinguish this PAMP-like T4 platform from other subunit vaccine platforms and adjuvant systems^28, 29, 48, 49^.

Ninth, the T4 nanoparticle vaccine does not require any adjuvant to stimulate robust anti-CoV-2 immune responses, as demonstrated in two animal models. In addition to reducing cost and manufacturing complexity, the adjuvant-less T4 vaccine formulations might provide a safe alternative because reactogenicity to vaccines is often associated with chemical adjuvants used in traditional vaccine formulations. In numerous immunizations with T4 phage, in a variety of animal models including mouse, rat, rabbit, and macaque, no significant adverse reactions were noted^28, 48–50^.

Finally, our best T4 vaccine candidate elicited, in addition to spike-specific antibodies, broad antibody responses against two additional virion components, E that is exposed on the surface and NP that is abundantly present on SARS-CoV-2 infected host cells^32, 33^. Furthermore, the spycatcher phage could serve as a “plug-and play” backbone to capture ectodomain trimers from other coronaviruses such as SARS-CoV-1 and MERS^2^, or any spytagged antigen(s) from other infectious agents. Thus, it is conceivable that different vaccine formulations can be “created” at the site of administration by mixing the spycatcher T4 phage with the desired antigen(s) combinations. Co-carrying multiple and distinct antigens in the same formulation increase the breadth of immune responses^51, 52^ and since regions of S-ecto, Ee, and NP are conserved in other coronaviruses^1, 2^, the T4 vaccine platform might be considered for expanded protection.

In conclusion, we have developed a versatile “universal” nanovaccine design template by which vaccine candidates can be rapidly generated, evaluated in animal models, and transitioned to human clinical trials as is being done currently with the T4-CoV-2 vaccine. As T4 is a highly stable nanoparticle, has good safety profile, and can be manufactured at a relatively low cost, it provides a robust platform to rapidly generate effective vaccines against any epidemic or pandemic pathogen in the future, particularly when multivalent vaccines are essential to protect global communities.

## Acknowledgments

We thank Dr. Victor Padilla-Sanchez and Frontera Supercomputer in Texas for assistance with the preparation of the video and Figure 1, Drs. Zhiqing Wang and Thomas Close at Purdue University for the Cryo-electron microscopy images, Drs. Barney Graham and Kizzmekia S. Corbett, (National Institutes of Health), and Jason S. McLellan (University of Texas, Austin) for providing recombinant plasmids containing SARS-CoV-2 Spike (S) genes, and Dr. Rayavara Kempaiah (UTMB) for providing the inoculum of the mouse-adapted SARS-CoV2 virus.

Funding: This research was supported by NIAID/NIH supplement grant 3R01AI095366-07S1 (subaward: 1100992-100) and in part by NIAID/NIH grants AI111538 and AI081726 and National Science Foundation grant MCB-0923873 to V.B.R. ViroVax LLC gratefully acknowledges the NIAID/NIH Supplement contract HHSN272201800049C to S.A.D. Special funding provided by the IHII-COVID19 pilot grant to A.K.C. is greatly acknowledged.

## Author contributions

V.B.R. designed and directed the project. J.Z. and V.B.R. designed research. J.Z., N.A., S.J., H.B., W-C.T. conducted most of the experiments. D.A.L, M.L.R., and S.A.D. performed the rabbit immunization and neutralization antibody titer assay. A.D. performed neutralization assay using the live virus with immune sera derived from mice. A.K.C. conducted mice challenge and neutralization titer assay. J.Z. and V.B.R. analyzed and interpreted the data. V.B.R. and J.Z. wrote the manuscript.

## Methods

### DNA, bacteria, and bacteriophages

The expression vector pET28b (Novagen, MA) was used for donor plasmid construction and protein expression plasmid construction. The spacer plasmids LbCpf1 and SpCas9 were constructed as described previously^21, 22^. The DNA fragments containing NP and RBD were codon-optimized for *E.coli* expression and synthesized by GeneArt (Thermo Fisher). The plasmids containing wild-type (WT) SARS-CoV-2 Spike (S) gene and S-ecto-6P gene were provided by Drs. Barney Graham, and Kizzmekia S. Corbett (National Institutes of Health), and Dr. Jason S. McLellan (University of Texas, Austin), respectively. The RBD gene for mammalian expression was amplified from the WT Spike (S) gene. SpyCatcher/Spy-tag and SUMO containing plasmids were purchased from Addgene (#133449 and #111560).

*E. coli* DH5α [*hsdR17 (rK–mK+) sup^2^*] (NEB) was used for all the clone constructions. The *E. coli* BL21-CodonPlus (DE3)-RIPL (Novagen, MA) was used for the expression of recombinant proteins. *E. coli* P301 (*sup*^*0*^) and *E. coli* B834 (*hsdRB hsdMB met thi sup^0^*) were used for the propagation and recombination of phages without amber mutations. The *E. coli* BL21 (DE3) RIPL transformed with amber-suppressor-1 plasmid and *E. coli* B40 (*sup*^*1*^) were used for propagation and recombination of phages with amber mutations (CTS-amber-NP). WT T4 phage was propagated on *E.coli* P301 or B834 and used as a starting phage for CRISPR engineering.

### Plasmid construction

CRISPR-LbCpf1/SpCas9 plasmid was constructed based on the streptomycin-resistant plasmid DS-SPCas (Addgene no. 48645) as described previously^21, 22^. The spacer containing fragments were prepared by annealing and extension of two amplified DNA fragments containing 26 bp complementary nucleotides (overlap extension PCR). The spacer fragment digested with restriction enzymes XhoI and EagI was cloned into linearized LbCpf1/SpCas9 plasmid. Sequences of the spacers are shown in Supplementary Table.

The donor plasmids for deletion/insertion, including Hoc-del, Soc-del, FarP7K-del, FarP18K-del, 39-56 11K-del, SegF-del, IPIII-del, IPII-del, E insertion (Hoc site), Soc-SpyCatcher/RBD insertion (Soc site), CTS-amber-NP insertion (IPIII site), CAG-S-fl/S-ecto insertion (FarP7K site), and CMV-RBD insertion (SegF site), were constructed using overlap extension PCR. Briefly, the DNAs including ~500 bp homology arms (left and right) with a desired deletion in the middle were amplified from T4 genome DNA, stitched, and cloned into pET28b linearized with BglII and XhoI to generate the deletion donor plasmid. For the insertion donor plasmid, the insertion fragment, left homology arm, and right homology arm, which contains 25 bp complementary nucleotides were stitched by annealing and overlap extension. The BglII and XhoI digested donor fragment was cloned into pET28b plasmid.

The Soc-fusion expression plasmids, including Soc-SpyCatcher, Soc-RBD, RBD-Soc, SUMO-Soc-Spy, and Soc-truncated RBDs (RBD67, RBD106, RBD135, RBD162, RBD181, and RBD197), were constructed by two rounds of cloning. First, the multiple cloning site (MCS) (NcoI, NdeI, NheI, and BmtI)-linker (4GGS)-Soc-linker (2GGGGS)-MCS (HindIII, EagI, NotI, and XhoI) was amplified using Soc-plasmid template and cloned into the pET28b DNA linearized with NcoI and XhoI restriction enzymes to generate pET28b-MCS-L-Soc-L-MCS. Second, DNAs corresponding to SpyCatcher, SUMO, Spy, and various RBDs were amplified and inserted to 3’- or 5’-MCS of pET28b-MCS-L-Soc-L-MCS as needed. CTS-NP and Ee-Hoc expression fragments were amplified using synthesized NP and T4 genomic DNA respectively, and cloned into NcoI and XhoI-linearized pET28b.

The plasmids for expression in mammalian cells, including pCMV-CD5-RBD, pCAG-CD5-S-fl, pCAG-CD5-S-ecto-6P, and pCAG-CD5-S-ecto-6P-Spytag, were constructed using standard protocols. The RBD fragment was amplified using the WT spike gene, and CD5 secretion leading peptide (MPMGSLQPLATLYLLGMLVASVLA) was added to the N terminus of RBD by PCR. The CD5-RBD was directionally cloned into pAAV vector (Cell Biolabs) using the HindIII and XhoI restriction enzyme sites (pCMV-CD5-RBD). For the construction of pCAG-CD5-S-fl, pCAG-CD5-S-ecto-6P, and pCAG-CD5-S-ecto-6P-Spytag, plasmids pCAG-S-fl and pCAG-S-ecto-6P were used as template and backbone. The CD5 fragment was cloned into N terminus of S-fl or S-ecto-6P using KpnI and EcoRI restriction sites. Similarly, Spytag (RGVPHIVMVDAYKRYK) was cloned into the C terminus of S-ecto using BamHI and XhoI restriction sites. All the constructed plasmids were sequenced to confirm 100% accuracy of the recombinant fragment (Retrogen, CA).

### Plaque assay

Plaque assays were performed to determine the efficiency of the individual spacers to restrict T4 phage infection. The CRISPR-Cpf1/Cas9 spacer plasmid was transformed into *E. coli* strains B834 or B40. Serially diluted T4 phages, in the range of 10^1^ to 10^7^ plaque forming units (pfu) in 100 μl Pi-Mg buffer (26 mM Na_2_HPO_4_, 68 mM NaCl, 22 mM KH_2_PO_4_, 1 mM MgSO_4_, pH 7.5), were mixed with 350 μl of spacer-containing *E. coli* (~10^8^ cells/ml). *E. coli* cells without spacer were used as a control. After incubation at 37°C for 7 min, 3.5 ml of 0.75% top agar with spectinomycin (50 μg/mL) was added to each tube, mixed, and poured onto LB plate. The plates were incubated at 37°C overnight to allow the formation of plaques. The pfu were counted on each plate and the efficiency of plating (EOP) was determined by dividing the pfu produced from infection of *E. coli* containing a spacer with the input pfu.

### CRISPR-mediated phage T4 genome editing and recombination

The CRISPR-Cpf1/Cas9 spacer plasmid and the corresponding donor plasmid were co-transformed into *E. coli* strains, either B834/P301 without amber suppressor, or B40/RIPL with amber suppressor. *E.coli* cells transformed with single plasmids, either the donor plasmid or the CRISPR spacer plasmid, were used as controls. An appropriate amount of T4 phages, as determined by the EOP as described above, were added to *E. coli* and incubated for 7 min at 37°C. After adding 3.5 ml of 0.75% top agar containing 50 μg/ml spectinomycin and 50 μg/ml kanamycin, the infection mixture was poured onto LB plate and incubated overnight. Single plaques, namely Generation 1 (G1), were picked using a sterile Pasteur glass pipet and transferred into a 1.5 ml Eppendorf tube containing 200 μl of Pi-Mg buffer. After 20 min incubation at room temperature with gentle vortexing every 5 min, serially diluted G1 phages were used to infect spacer-containing *E. coli* cells (50 μg/ml spectinomycin). This eliminated any parental phage background under CRISPR pressure. The resultant single G2 plaques were picked and used to infect *E. coli* cells (without spacer or donor) to produce purified G3 phages. Single G3 plaques were then picked into 200 μl of Pi-Mg buffer. PCR analysis was performed to confirm DNA deletion or foreign gene insertion. One microliter of G3 phages were denatured at 94°C for 8 min and used as a template for PCR using Phusion High-Fidelity PCR Master Mix (Thermo Fisher). The amplified DNA fragment was purified using QIAquick Gel Extraction Kit (Qiagen) and sequenced (Retrogene). The G3 phages in which the recombinant DNA sequence was confirmed with 100% accuracy were selected and a few drops of chloroform were added. These plaque-purified “zero stocks” were stored at 4°C. More rounds of CRISPR gene editing were similarly introduced into the same phage as needed as described above.

### Phage production and purification

*E. coli* strains B40 or B834 were used for the production of amber-phage or non-amber-phage, respectively. Fresh overnight *E. coli* cells were inoculated in 1 L of Moore’s media (20 g tryptone, 15 g yeast extract powder, 2 g dextrose, 8 g NaCl, 2 g Na_2_HPO_4_, 1 g KH_2_PO_4_, dissolved in 1 L MQ water) at 1/50 dilution, and then grown at 37°C for 2-2.5 hr in a shaker incubator at 200 RPM. When the cells reached a density of ~4 × 10^8^/ml, phages were added at a multiplicity of infection (MOI) of 0.5. The infection mixture was cultured at 37°C, 200 RPM, for another 2.5-3 hr and periodically checked under the microscope. As the cells get infected with phage, the shape of cells change from long bacilli to short dumbbell shape. When the cell number started dropping, the culture was transferred to Sorvall GSA bottle and centrifuged at 27, 504 g for 1 hr at 4 °C. The supernatant was discarded, and the pellet was resuspended in 50 ml Pi-Mg buffer containing 10 μg/ml DNase I (or Benzonase) and one tablet of protease inhibitor cocktail. The resuspended pellet was added with 5 ml chloroform and incubated at 37°C for 1 hr to lyse the bacteria and release the phage. The debris was removed by low-speed centrifugation at 4,302 g for 10 min. The phage-containing supernatant was transferred to a sterile falcon tube and the pellet was discarded. The titer of this phage stock was determined using *E. coli* B40 or B834. The working stock of phages can be stored at 4°C for long periods of time with a few drops of chloroform added to prevent microbial contamination, or used as seed phage to make SpyCatcher or Soc-RBD displayed phage, or purified as a vaccine candidate for animal studies.

For purification of phage, ~50 ml phage stock was distributed into 5 Corex glass tubes (10 ml each) and sealed with parafilm. The phage stock was centrifuged at 34,540 g for 1 hr using a Sorvall SS34 rotor. The supernatant was discarded and the phage-containing pellet was resuspended with 1-3 ml Pi-Mg buffer plus 5 μl Benzonase overnight at 4°C. Next, the resuspended phages were loaded onto a CsCl gradient and centrifuged at 152,000 g for 1 hr in a Beckman ultracentrifuge using a SW 55Ti swinging bucket rotor. The purified phage band localized between CsCl densities 1.46 and 1.55 was collected by inserting a syringe right below the phage band and aspirating the band. The phages were dialyzed in high-salt Tris-Mg buffer (10 mM Tris-HCl, pH 7.5, 200 mM NaCl, 5 mM MgCl_2_) for 4 hr followed by low-salt Tris-Mg buffer (10 mM Tris-HCl, pH 7.5, 50 mM NaCl, 5 mM MgCl_2_) overnight. The second-round CsCl centrifugation and dialysis of purified phages were applied to obtain purer phages. Two-round-CsCl-purified phages were further purified by passing through a 0.22 μm filter to remove any minor bacterial contaminants. The phage concentration and copy numbers of displayed antigens were quantified by 4-20% SDS–polyacrylamide gel electrophoresis (SDS-PAGE).

### Production of Soc-SpyCatcher and Soc-RBD displayed phages

*E.coli* BL21 (DE3) RIPL transformed with T7-Soc-Spycatcher (or Soc-RBD) plasmid were used for Soc-Spycatcher (or Soc-RBD) *in vivo*-displayed phage production. *E.coli* BL21 (DE3) RIPL co-transformed with T7-Soc-Spycatcher (or Soc-RBD) plasmid and amber suppressor plasmid was used for the production of Soc-Spycatcher (or Soc-RBD) displayed and NP protein packaged phage. Briefly, the RIPL cells were inoculated into 1 L of Moore’s Media at 1/50 dilution with appropriate antibiotics (RIPL-Soc-Spycatcher: 50 μg/ml Kanamycin+ 37 μg/ml Chloramphenicol; RIPL-Soc-Spycatcher-Amber Suppressor: 50 μg/ml Kanamycin + 37 μg/ml Chloramphenicol + 100 μg/ml Ampicillin). The culture was incubated at 37°C, 200 RPM, for 2.5-3 hr. When the cells reached a density of ~4 × 10^8^/ml, 0.5 mM IPTG was added for induction of recombinant protein expression. At 10 min post IPTG addition, the corresponding phages (Hoc-del/Soc-del/IPII-del/IPIII-del) were added to infect cells at an MOI of 0.5. The culture was further incubated at 37°C, 200RPM, for 3 hr. The following phage production and purification procedures are the same as described above.

### S-trimer expression and purification

Plasmid pCAG-CD5-S-ecto-6P-Spytag was transiently transfected into ExpiCHO cells using ExpiFectamine CHO transfection kit (Thermo Fisher). After 18-22 hr of transfection, cells were supplemented with ExpiCHO Feed and Enhancer and grown at 32°C according to the manufacturer’s High Titer protocol. Cultures were harvested 8-10 days after transfection by centrifuging the cells at 3000 g for 20 min at 4°C. The supernatant was clarified through a 0.22 μm filter and then loaded on a HisTrap HP column (Cytiva) previously equilibrated with wash buffer (50 mM Tris-HCl, pH 8.0, containing 300 mM NaCl and 20 mM imidazole), at a flow rate of 1 ml/min, using AKTA Prime-Plus liquid chromatography system (GE Healthcare). Protein-bound column was washed with wash buffer until the UV absorbance reached the baseline to remove non-specifically bound proteins. The trimers were eluted using a 20 mM-300 mM linear gradient of imidazole. HisTRAP eluted peak fractions were pooled and applied to a Hi-Load 16/600 Superdex-200 (preparation grade) size-exclusion column (GE Healthcare) equilibrated with the gel filtration buffer (50 mM Tris-HCl, pH 8, 150 mM NaCl) to obtain purified trimers, using the ÄKTA FPLC system. Eluted fractions were collected, filtered through 0.22 μm filter unit, flash-frozen, and stored at −80 °C until use.

### Quantification of S-trimer and rRBD display on T4-SpyCatcher phage

*In vitro* display of S-trimer/rRBD on the T4-SpyCatcher phage was assessed by co-sedimentation as described previously with some modifications^27, 28^. Briefly, two-round CsCl purified and 0.22 μm filtered phage particles were sedimented for 45 min at 34,000 g in Protein-LoBind Eppendorf tubes, washed twice with sterilized phosphate-buffered saline (PBS) buffer (pH 7.4), and resuspended in PBS buffer (pH 7.4). S-trimer/rRBD was sedimented for 25 min at 34,000 g to remove any possible aggregates present in the sample. T4-SpyCatcher phages were incubated with S-trimer/rRBD proteins at 4°C for 1 hr. The mixtures were sedimented by centrifugation at 34,000 g for 45 min, and unbound protein in the supernatants was removed. After washing twice with excess PBS to further remove the unbound protein and any other minor contaminants, the phage pellets containing the displayed proteins were incubated at 4°C overnight and then resuspended in PBS. For rabbit animal studies, fifty microliters of phage-trimer particles were added to blood agar (TSA with sheep blood) to examine any contamination of a wide variety of fastidious microorganisms. The resuspended pellets were analyzed using Novex 4-20% SDS– SDS-PAGE mini gel (Thermo Fisher Scientific, Waltham, MA) to quantify the S trimer/rRBD copies. After Coomassie Blue R-250 (Bio-Rad, CA) staining and destaining, the protein bands on SDS-PAGE gels were scanned and quantified by ChemiDoc MP imaging system (BioRad) and image J. The copy numbers of SpyCatcher and displayed S-trimer/rRBD molecules per capsid were calculated using gp23 (major capsid protein; 930 copies) or gp18 (major tail sheath protein; 138 copies) as internal controls and S-trimer protein standard.

### SUMO-RBD-spy protein expression and purification

The *E.coli* expression, denaturation, refolding, and purification of SUMO-RBD-Spy (rRBD) were performed using a similar procedure described previously^53^. Briefly, the BL21-CodonPlus (DE3)-RIPL cells containing PET28b-SUMO-RBD-Spy expression plasmid were induced with 0.5 mM isopropyl--D-1-thiogalactopyranoside (IPTG) for 3 h at 28°C. Cells were harvested and resuspended in buffer A (20 mM Tris–HCl, 500 mM NaCl, 5 mM imidazole, 5 mM β-mercaptoethanol, pH 7.9) containing protease inhibitor cocktail (Roche, USA, Indianapolis, IN) and Benzonase. After the cells were lysed using a French press (Aminco, Urbana, IL) and centrifuged, the pellet containing the inclusion body proteins was resuspended and washed with buffer B (buffer A + 0.5% Triton X-100). Then, the inclusion bodies were solubilized in buffer C (Buffer A + 8 M urea) by incubating/stirring at 4 °C overnight, followed by centrifugation and clarification. The supernatant containing the denaturing protein was loaded on HisTrap column (AKTA-prime; GE Healthcare) followed by washing with buffer C. The rRBD was refolded on HisTrap column using a linear gradient of 8 to 0 M urea containing buffer D (20 mM Tris–HCl, 500 mM NaCl, 5 mM imidazole, 1 mM GSH, 0.1 mM GSSG, 20% glycerol, pH 7.9). Finally, the column was washed with buffer E (20 mM Tris–HCl, 500 mM NaCl, 100 mM imidazole, 20% glycerol, 5% glucose, pH 7.9), and the refolded rRBD was eluted with buffer F (20 mM Tris–HCl, 500 mM NaCl, 800 mM imidazole, 20% glycerol, 5% glucose, pH 7.9) and dialyzed to remove imidazole. The proteins were then quantified, aliquoted, and stored at −80°C until use.

### Western blot analysis

After treatment with multiple freeze-thaw cycles and Benzonase, phage particles were boiled in SDS loading buffer for 10 min, separated by 4-20% SDS-PAGE, and transferred to nitrocellulose membrane PVDF (Bio-Rad). The PVDF was then blocked with 5% bovine serum albumin (BSA)-PBS (pH 7.4) buffer at RT for 1 hr with gentle shaking. Anti-NP or anti-RBD primary antibodies were added to the blots and incubated overnight at 4°C in PBS-5% BSA, followed by five times rinsing in PBST buffer [1×PBS (pH 7.4) and 0.05% Tween 20]. Goat-anti-mouse or goat-anti-rabbit HRP-conjugated antibody (Thermo Fisher) was applied at a 1:5000 dilution in 5% BSA-PBST for 1 hr at RT with gentle shaking. After rinsing five times in PBST, binding was visualized with an enhanced chemiluminescence substrate (Bio-Rad) using the Bio-Rad Gel Doc XR+ System and Image Lab software according to the manufacturer’s instructions (Bio-Rad).

### Cell culture and transfection

HEK293T cells were maintained in Dulbecco’s modified Eagle’s medium (DMEM; Gibco) supplemented with 1% antibiotics (Thermo Fisher), 1x HEPES (Thermo Fisher), and 10% fetal bovine serum (Thermo Fisher). Cells were passaged with 0.25% (w/v) trypsin/0.53 mM EDTA at a sub-cultivation ratio of 1:5 at 80 to 90% confluence. Cultures were incubated in a humidified atmosphere at 37°C and 5% CO2. The recombinant plasmid containing human ACE2 gene (Addgene #1786) was transfected into HEK293T cells using Lipofectamine 2000 Transfection Reagent (Thermo Fisher) according to the manufacturer’s instructions. Two days after ACE2 plasmid transfection, the cells were used for RBD or phage binding assay.

### Inhibition by mice sera of RBD binding to cell surface ACE2

Human ACE2 transfected HEK293T cells were washed with PBS twice and then fixed with 4% formaldehyde for 15 min at RT. After rinsing twice in PBS for 5 min each, cells were blocked in blocking buffer (5% BSA-PBS) for 1 hr at RT. Recombinant SARS-CoV-2 RBD protein (Sino Biological) was added to the cells to a final concentration of 0.2 μg/ml in the presence or absence of the sera with a series of dilutions. The unbound RBD was removed by washing cells five times with PBST (PBS + 0.1% Tween 20) for 5 min each. The 1/1000 diluted human anti-RBD monoclonal antibody (Thermo Fisher) was added to cells and incubated in a humidified chamber for 1 hr at RT or overnight at 4°C. After rinsing five times in PBST for 5 min each, Alexa488- or Rhodamine-conjugated goat anti-human secondary antibody was added (1/500 dilution) (Thermo Fisher), and incubated for 2~3 hr at room temperature in the dark. The cells were then rinsed five times in PBST for 5 min each and counter-stained with 1 μg/ml Hoechst 33342 (Thermo Fisher) for 5 min. The fluorescent signals were recorded by fluorescence microscopy (Carl Zeiss).

### Binding of T4-S trimer-GFP phages to cell surface ACE2

Soc-GFP protein was produced as described previously^27^. *In vitro* display of Soc-GFP on the T4-SpyCatcher or T4-S trimer phage was assessed by the co-sedimentation and SDS-PAGE, similar to the procedures of S-trimers displayed on T4 phage. The T4-SpyCatcher-GFP or T4-S trimer-GFP phages were resuspended in Opti-MEM medium (Thermo Fisher) and then added to ACE2-transfected HEK293T cells. After 6 hr incubation, the unbound phages were removed by rinsing three times in PBS for 5 min each. The GFP fluorescence was recorded by fluorescence microscopy (Carl Zeiss).

### Mouse immunizations

We followed the recommendations of the National Institutes of Health about mouse study (*the Guide for the Care and Use of Laboratory Animals*). All mouse experiments were approved by the Institutional Animal Care and Use Committee of the Catholic University of America (Washington, DC) (Office of Laboratory Animal Welfare assurance number A4431-01) and the University of Texas Medical Branch (Galveston, TX) (Office of Laboratory Animal Welfare assurance number A3314-01). The SARS-CoV-2 virus challenge study was conducted in the animal biosafety level 3 (ABSL3) suite.

Six- to eight-week-old female BALB/c mice (The Jackson Laboratory) were randomly grouped (5 mice per group) and allowed to acclimate for 14 days. The phage vaccine candidates were administered by intramuscular (i.m.) injections into their hind legs. A total of 6 ×10^11^ phages carrying approximately 20 μg of SARS-CoV-2 antigen(s) were injected on days 0 (prime), 21 (boost 1), and 42 (boost 2). Negative control mice received the same volume of PBS buffer (Naive) or the same amount of T4 control phage. A group of mice immunized with purified S trimer (20 μg) adjuvanted with Alhydrogel was included as the positive control. Blood was drawn from each animal on days 0 (pre-bleed), 14, 35, and 56, and the isolated sera were stored at −80°C.

### Rabbit immunizations

All experiments were performed at Envigo/Cocalico Biologicals (Reamstown, PA) in accordance with institutional guidelines. Adult New Zealand White rabbits were immunized intramuscularly in the flank region with 3X10^11^ PFU T4 phages/dose in 0.2 mL saline (n = 4 for group). Pre-immune test-bleeds were first obtained via venipuncture of the marginal vein of the ear on Day 1. Animals were immunized on Days 1, and 15 (Prime + One-boost regimen). Immune sera were obtained on Day 25.

### ELISA determination of IgG and IgG subtype antibodies

ELISA plates (Evergreen Scientific) were coated with 100 μl per well of 1 μg/ml of SARS-CoV-2 S-ecto protein (Sino Biological), SARS-CoV-2 RBD-untagged protein (Sino Biological), SARS-CoV-2 NP protein (Sino Biological), or SARS-CoV-2 E protein (1-75 aa) (Thermo Fisher) in coating buffer [0.05 M sodium carbonate–sodium bicarbonate (pH 9.6)]. After overnight incubation at 4°C, the plates were washed twice with PBS buffer and blocked for 2 hr at 37°C with 200 μl per well PBS–5% BSA buffer. Serum samples were diluted with a 5-fold dilution series beginning with an initial 100-fold dilution in PBS–1% BSA. One hundred microliters of diluted serum samples were added to each well and the plates were incubated at 37°C for 1 hr. After washing five times with PBST (PBS + 0.05% Tween-20), the secondary antibody was added at 1:10,000 dilution in PBS–1% BSA buffer (100 μl per well) using either goat-anti-mouse IgG-HRP, goat-anti-mouse IgG1-HRP, goat-anti-mouse IgG2a-HRP (Thermo Fisher), or goat-anti-rabbit IgG-HRP (Abcam). After incubation for 1 hr at 37°C and five washes with PBS-T buffer, plates were developed using the TMB (3,3’,5,5’-tetramethylbenzidine) Microwell Peroxidase Substrate System (KPL). After 5-10 min, the enzymatic reaction was stopped by adding TMB BlueSTOP (KPL) solution. The absorbance was read within 30 min at 650 nm on a VersaMax spectrophotometer. The endpoint titer was defined as the highest reciprocal dilution of serum that gives an absorbance more than 2-fold of the mean background of the assay.

### Binding of T4 displayed RBD or S-trimer to human ACE2 protein

An ELISA to analyze the binding of RBD, S-ecto-6P-spytag trimer, T4 displayed RBD/S-trimer to human ACE2 protein was performed as described above. Briefly, 100 ng protein or 1×10^10^ phages were coated on plates overnight at 4°C. After blocking with PBS–5% BSA buffer, recombinant human ACE2-mouse Fc protein (Sino Biological) with a series of dilution was added and incubated for 1 hr at 37°C. Plates were then incubated with the secondary goat-anti-mouse IgG-HRP antibody and developed with TMB substrate. Reactions were stopped and the absorbance was measured at 650 nm on a VersaMax spectrophotometer.

### Virus Neutralization assay using BSL-3 live SARS-CoV-2

Neutralizing antibody titers in mouse immune sera were quantified by Vero E6 cell-based micro-neutralization assay using SARS-CoV-2_US-WA-1/2020 strain as previously described^46^. Briefly, serially 1:3 downward diluted mouse sera that were decomplemented at 56°C for 60 min in 60 μl volume were incubated for 1 h at room temperature in duplicate wells of 96-well microtiter plates that contained 120 infectious SARS-CoV-2 virus in 60 μl in each well. After incubation under BSL-3 conditions, 100 μl of the mixtures in individual wells were transferred to Vero E6 cell monolayer grown in 96-well microtiter plates that containing 100 μl of MEM/2%FCS medium in each well and were cultured for 72 h at 37°C before assessing the presence or absence of cytopathic effect (CPE). Neutralizing antibody titers of the tested specimens were calculated as the reciprocal of the highest dilution of sera that completely inhibited virus-induced CPE in at least 50% of the wells and expressed as 50% neutralizing titer (NT_50_).

Neutralizing antibody titers in rabbit immune sera were quantified using an automated, liquid-handler-assisted, high-throughput, microfocus neutralization/high-content imaging methods developed at ViroVax. Briefly, rabbit sera (paired pre-immune and immune), were first decomplemented at 56°C for 60 min, and were then serially diluted in 384 well plates, in duplicate, using a BioTek Precision 2000 liquid handler, along with two reference sera. Twenty microliter aliquots of SARS-CoV-2_USA-WA1/2020 were added to all test wells and positive control wells to yield a final MOI of 10 under BSL-3 conditions. Vero (ATCC® CCL-81) cells were maintained in a high-glucose Dulbecco’s modified Eagle’s medium (DMEM) supplemented with 10% fetal bovine serum (FBS; HyClone Laboratories, South Logan, UT) and 1% penicillin/streptomycin at 37°C with 5% CO2. After preincubating the plates for 1 hr, 20 μl of Vero cells (10^6^/ml), containing 50 μg/mL of propidium iodide (PI), was added to all wells using the liquid handler. Plates were then loaded in an IncuCyte S3 high-content imaging system (Essen Bioscience/Sartorius, Ann Arbor, MI). Longitudinal image acquisition and processing for virus-induced cytopathic effect (CPE) and cell death (PI uptake) were performed every six hrs, until cell death profiles had crested and stabilized (3.5 days). Neutralizing antibody titers (expressed as IC50 or IC90) were obtained from four-parameter logistic curve-fits of cell death profiles using OriginPro 9 (Origin Lab Corp., Northampton, MA).

### Challenge of the mice with mouse-adapted live BSL-3 SARS-CoV-2 virus

Immunized mice were challenged with the mouse-adapted (MA) SARS-CoV-2/MA10 strain^47^, a generous gift from Ralph Baric at UNC, by the intranasal route as previously described^54^. Briefly, mice were inoculated with 60 μl of SARS-CoV2-MA10 at a dose of ~10^5^ TCID50. The animals were weighed every day over the indicated period of time for monitoring the onset of morbidity (weight loss and other signs of illness) and mortality, as the endpoints for evaluating the vaccine efficacy.

### Statistics

All the data were presented as means ± SEM except where indicated. Statistical analyses were performed by Graph Pad Prism 9.0 software using one-way or two-way Analysis of variance (ANOVA) according to the data. Tukey’s multiple comparisons post-test was used to compare individual groups. Significant differences between two groups were indicated by *P < 0.05, **P < 0.01, ***P < 0.001, and ****P < 0.0001. ns, no significance. P-values of < 0.05 were considered significant.

---

**Supplementary Table.**
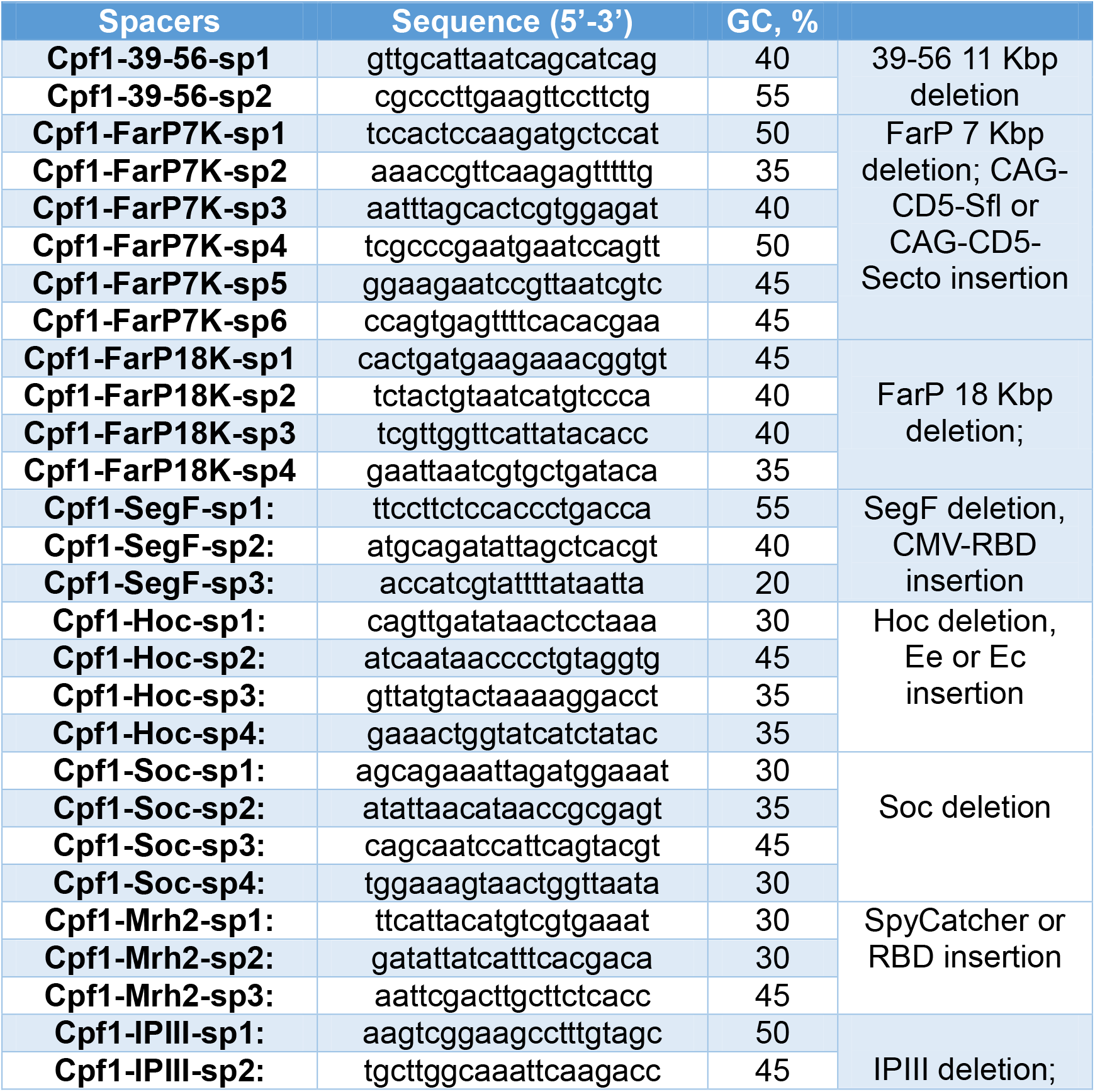
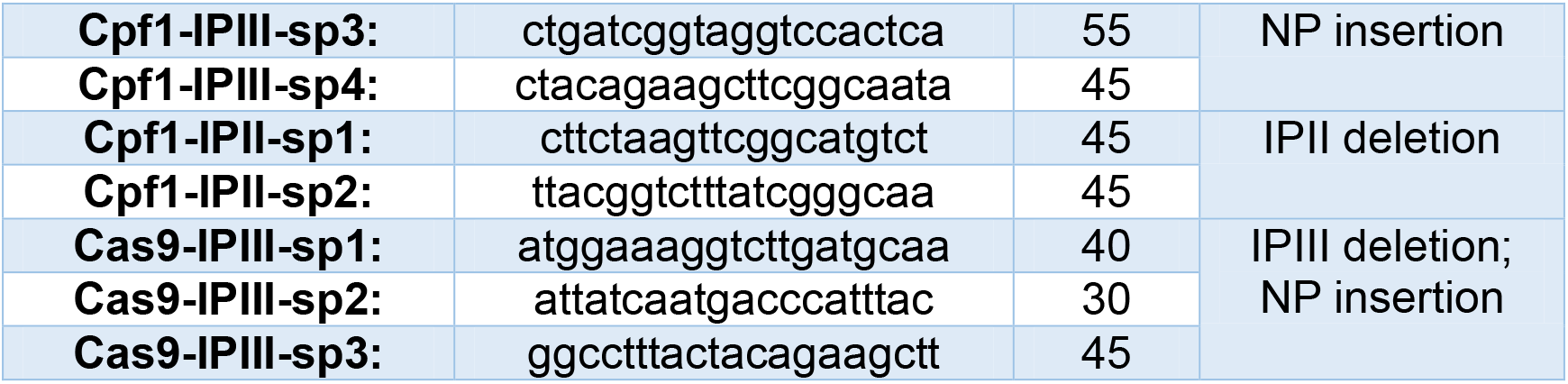
Cpf1 spacer information used in this study.

